# Anatomically precise relationship between specific amygdala connections and selective markers of mental well-being in humans

**DOI:** 10.1101/2020.03.08.980995

**Authors:** Miriam C Klein-Flügge, Daria EA Jensen, Yu Takagi, Lennart Verhagen, Stephen M Smith, Matthew FS Rushworth

## Abstract

There has been increasing interest in using neuroimaging measures to predict psychiatric disorders. However, predictions usually rely on large numbers of brain connections and large disorder heterogeneity, thus lacking both anatomical and behavioural specificity, preventing the advancement of targeted interventions. Here, we address both challenges. First, using resting-state functional MRI, we parcellated the amygdala, a region implicated in mood disorders but difficult to image with high fidelity, into seven nuclei. Next, a questionnaire factor analysis provided four sub-clinical latent behaviours frequently found in anxious-depressive individuals, such as negative emotions and sleep problems. Finally, for each latent behaviour, we identified the most predictive connections between individual amygdala nuclei and highly specific regions of interest e.g. dorsal raphe nucleus in the brainstem or medial prefrontal cortical regions. A small number of distinct connections predicted behaviours, providing unprecedented levels of specificity, in humans, for relating mental well-being to precise anatomical connections.

## Introduction

It has become increasingly popular, in recent years, to use measures derived *in vivo* from human magnetic resonance imaging (MRI) to predict health outcomes, including measures of mental well-being. For example, resting-state functional MRI (rs-fMRI) connectivity measures can predict whether a person suffers from, or will respond to treatment for, Generalized Anxiety Disorder (GAD), Major Depressive Disorder (MDD), and obsessive-compulsive disorders (OCD) ^1–5^. The prediction accuracies achieved in these types of studies are often impressive and typically reach values between 60-80%. Yet, in the large majority of cases, predictions rely on a large number of brain regions, networks or connections. Hence the impressive prediction accuracies come at the significant cost of reduced anatomical specificity.

Despite the critical importance of such studies for diagnosis and prognosis, a lack of anatomical specificity may be problematic when the aim is a mechanistic understanding of the disease to support targeted treatment interventions. Identification and characterization of specific circuits may be necessary for establishing the nature and variants of the illness and it may be critical for developing new treatments that involve manipulation of brain activity in specific circuits.

A second problem is that unsupervised decoding methods, although powerful, are often agnostic to anatomical priors. Yet a large body of evidence has established the roles of specific neurotransmitter systems and particular brain regions in mediating important functions implicated in mental health. Limbic structures that mediate emotional processing and their connections with prefrontal regions are consistently reported to play an important role and one key hub within this network is the amygdala ^6–11^. Removal or disruption of this region reduces fear and anxiety responses ^10, 12–14^. Positron-emission tomography in depressed patients shows abnormal metabolism in amygdala and connected subgenual prefrontal cortex ^7, 10, 15^. And the amygdala is one of the key regions for regulating and expressing emotions ^6, 10, 12, 16–18^. An aim in the current study was, therefore, to examine the degree to which it is possible to explain variance in mental well-being across humans, including social and emotional behaviour, in relation to the functional connectivity of identifiable neural circuits – those centred on the amygdala. The monosynaptic connections of the amygdala to specific cortical and subcortical regions have been established for some time in animal models including primates ^19^ and there is increasing knowledge of the behaviours mediated by amygdala interactions ^11, 13, 20, 21^.

If, however, a decision is taken to focus on a brain region such as the amygdala then a third problem arises. Many of the key brain areas with which it interacts are in the brainstem where it has been difficult to image activity. Moreover, such regions have very specific connections to particular sub-nuclei within the amygdala. Therefore, our first step was to parcellate the human amygdala into constituent functional sub-units. We took advantage of the high-quality data acquired as part of the human connectome project (HCP; ^22^). Using resting-state measures from 200 healthy participants, we reliably identified seven amygdala nuclei within each hemisphere. We also invested considerable effort in developing a refined data pre-processing pathway that focused on the removal of breathing related artefacts that allowed us to examine activity even in brainstem regions, several of which exhibit very specific interactions with particular amygdala subnuclei.

In tandem with improving anatomical specificity we also aimed to tackle another major problem in relating baseline neural measures to mental well-being. Namely, the disorders themselves are ill-defined and span a broad range of impairments which are not consistently present in all patients diagnosed with the same disorder ^23^ and which are partly overlapping between disorders. This may be another reason why a classifier trained to distinguish a depressed from a non-depressed person is likely to reveal a broad network of regions instead of mapping onto well-defined and anatomically interpretable brain circuits. If we are able to focus on specific rather than broad symptom categories, we may better be able to relate them to specific brain circuits. Because of the sample we examined, mental health varied on a sub-clinical scale. Nevertheless, we were able to define latent behaviours by applying a factor analysis to a large number of questionnaire scores which captured four aspects of mental well-being. In our final and most critical step, we selected the best predictors in terms of connections between amygdala nuclei and other brain regions for each latent measure of mental well-being. We showed that a few specific connections predicted a reliable portion of the variance in each latent behaviour. Our study provides the first evidence in a large pool of healthy participants that using an anatomically informed approach and a more sensitive characterization of the behavioural phenotypes related to mental well-being, we can identify a small set of brain connections that can be used to predict latent aspects of mental well-being.

## Results

### In vivo parcellation of the human amygdala into seven anatomically plausible nuclei

*Post-mortem* histological examination in humans and other species has established that the amygdala is composed of anatomically distinct nuclei. Our first aim, therefore, was to use rs-fMRI to provide the best possible *in vivo* parcellation of human amygdala into its nuclei. Previous work in humans *in vivo* has delineated two or three subdivisions within the amygdala (e.g. basolateral versus centromedial) ^24–27^. However, given the quality of the HCP data (e.g., improved sequences, 2mm isotropic resolution, 0.7s temporal resolution, ∼1h rs-fMRI per person ^28^), and as a result of the enhanced processing steps we took to remove physiological confound signals, we reasoned that we might reliably identify a more detailed pattern of anatomical organization within the amygdala.

We generated a group connectome using carefully pre-processed rs-fMRI data from a subset of 200 HCP participants. Additional pre-processing focussed on removal of physiological artefacts and led to gains in temporal signal-to-noise (tSNR) in amygdala and many of its projection targets and sources, e.g., medial temporal lobe areas, subgenual prefrontal cortex, and most prominently subcortical and brainstem structures (see Methods and **Supplementary Fig 1, A-B**). We did not include all 1206 HCP participants because these additional pre-processing steps required good quality physiological recordings of respiration and cardiac activity which were not available in the remaining participants. The resulting group connectome, containing the average functional connectivity between each pair of brain-ordinates, therefore provided high-fidelity connectivity estimates of otherwise difficult to image regions. This is, for example, illustrated by the average amygdalae to whole brain connectivity (**Fig 1A**), which, in line with previous work ^19^, highlights overall strong functional coupling between the amygdala and lateral temporal and temporal pole regions, ventral caudal medial frontal cortex (BA32 and BA25), thalamus, hypothalamus, and ventral striatum.

**Figure 1.**
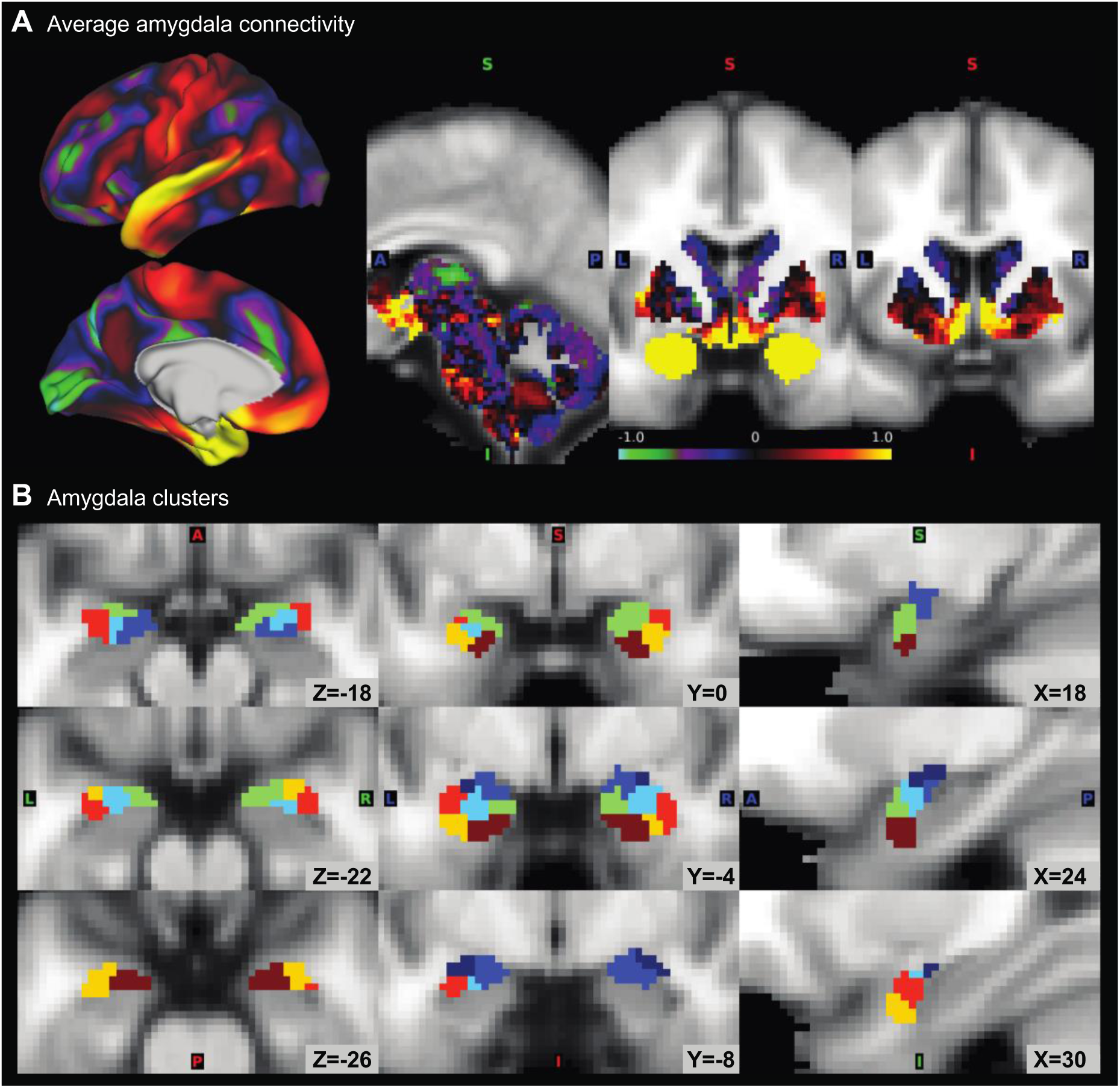
Average amygdala connectivity and definition of amygdala clusters. **A,** A group connectome was generated from resting-state fMRI (rs-fMRI) data of 200 HCP participants using an improved pre-processing pipeline to correct for physiological noise (Fig S1). The average functional coupling of all amygdala voxels to the rest of the brain, corrected for global absolute coupling strength, shows patterns that would be expected from tracer studies, for example strong connectivity of the amygdalae with subgenual ACC, hypothalamus, and ventral striatum. **B**, Hierarchical clustering was performed on the similarities between the whole-brain functional connectivity patterns of different amygdala voxels to identify amygdala subdivisions sharing connectivity profiles. Seven sub-divisions were identified (left: horizontal; middle: coronal; right: saggital view), showing strong symmetry across hemispheres and strong resemblance with subdivisions identified from histology and high-resolution *post-mortem* structural neuroimaging.

To identify subdivisions within the amygdala, hierarchical clustering was performed on the similarity of the whole-brain connectivity pattern between amygdala voxels. This resulted in parcellations of the amygdalae into increasing numbers of clusters. By carefully comparing the location and size of the obtained clusters to known anatomical subdivisions of the amygdala and using heuristics such as symmetry across hemispheres (see Methods), we chose a parsimonious and anatomically plausible parcellation for further analyses. This parcellation contained seven subdivisions in each hemisphere (**Fig 1B**; see **Supplementary Fig 1D** for other depths of clustering).

Several interesting features naturally emerged in this parcellation. First, clusters were nearly symmetrical across left and right hemispheres (**Fig 1B**). Importantly, this was not the consequence of constraints enforced by the clustering algorithm, and yet matched expectations from anatomical work because neurons with similar projection patterns tend to cluster in space, and inter-hemispheric similarities in the connection patterns of a given nucleus in each hemisphere outweigh their differences. Second, another naturally emerging feature of the data, again consistent with expectations from histological analysis, was that clusters were spatially cohesive but differed in size. For instance, a putative central nucleus contained 30 voxels per hemisphere, but the ventrolateral nucleus contained 50 voxels. Finally, the clusters were located in such a way that a clear progression from ventro-lateral to dorso-medial and from ventral-anterior to dorsal-posterior could be observed, thus corresponding to organizational principles reported previously (**Figure 1B**; ^29^).

To facilitate links to other studies, we assigned each cluster a putative label, corresponding to nuclei that have previously been identified (see Methods). As a guide, we used the best match in size and position when comparing our clusters with several atlases of the human amygdala ^29–31^ (**Fig 2A**). The seven nuclei were labelled central nucleus (Ce), cortical nuclei (CoN), auxiliary basal nucleus (AB), basal nucleus (B), and lateral nuclei (ventral portion: LaV, intermediate portion: LaI, dorsal portion: LaD).

**Figure 2.**
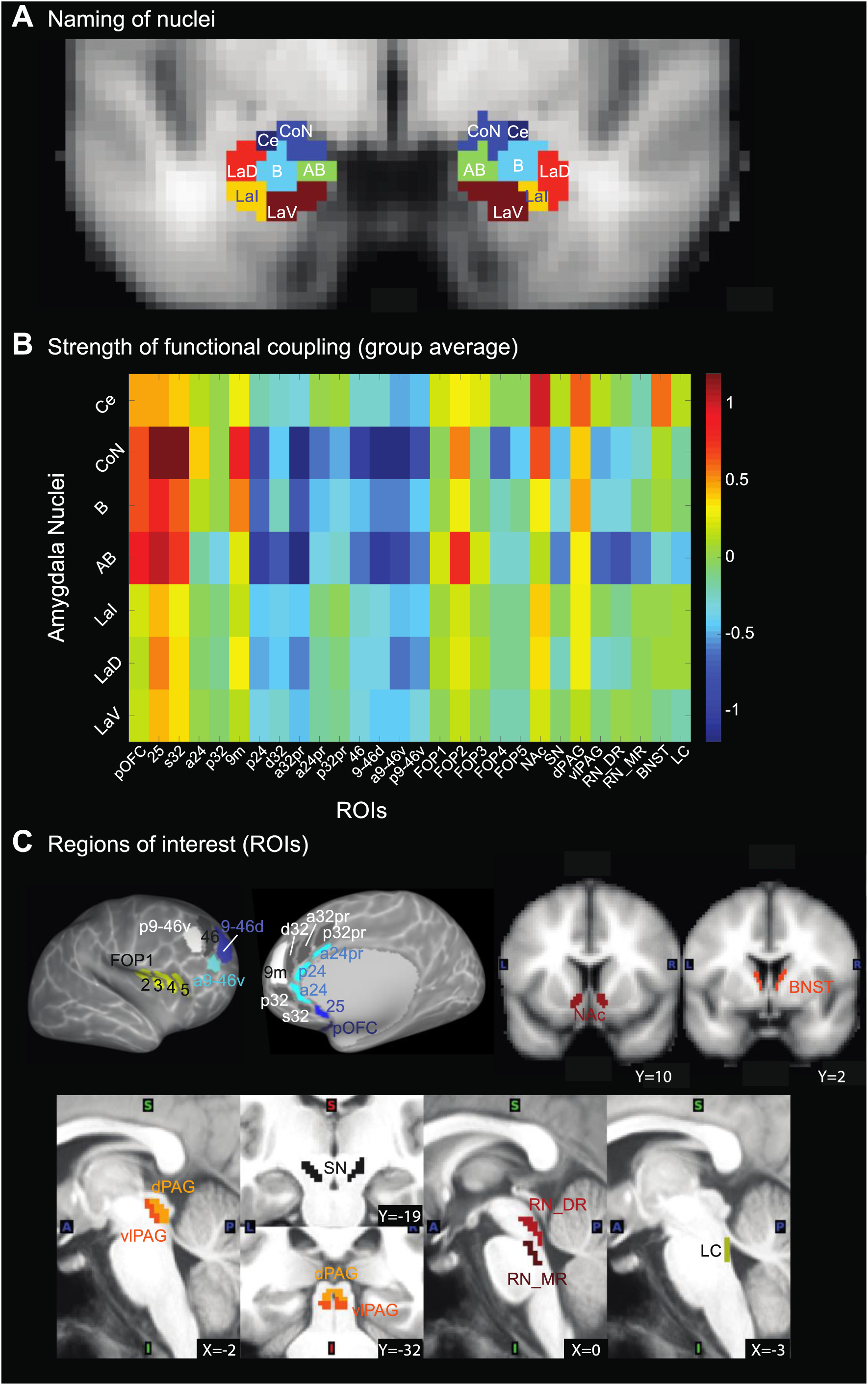
Amygdala nuclei and their profile of connectivity to regions of interest. **A,** Labels assigned to the seven amygdala subdivisions obtained from hierarchical clustering: Ce = central nucleus, CoN = cortical nuclei, B = basal, AB = auxiliary basal, LaV = lateral (ventral part), LaI = lateral (intermediate part), LaD = lateral (dorsal part). **B**, Average resting-state connectivity from the seven nuclei to 28 regions of interest (ROIs) defined *a priori* based on their potential role in regulating emotions and mental well-being. This highlights strong coupling of subgenual cortex (area 25) to the entire amygdala, but particularly to basal subdivisions, in line with tracer work. Similar profiles are observed for posterior OFC (pOFC) and the subgenual portion of area 32 (s32). By contrast, subcortical and brainstem regions most strongly connect with the central nucleus as expected. **C**, Masks of all ROIs used in this study. For details on their definition, please refer to the Methods. NAc=Nucleus Accumbens; BNST=bed nucleus of the stria terminalis; vl/dPAG=ventrolateral/dorsal periaqueductal grey; SN=substantia nigra; RN_DR/RN_MR=dorsal and median raphe nuclei; LC=locus coeruleus. Definitions of cortical regions were taken from Glasser et al., 2016.

One of the main aims of this study was to identify specific connections between amygdala nuclei and other brain regions that help regulate functions implicated in mental health variation (e.g. sleep and emotion variation). To identify regions of interest (ROIs) with which the amygdala interconnects, we therefore focussed on regions central to these processes (**Fig 2C**) and with known mono- or di-synaptic connectivity with the amygdala. In the brainstem, we defined ROIs in locus coeruleus (LC), dorsal and median raphe nuclei (DRN, MRN), dorsal and ventrolateral periaqueductal grey (dPAG, vlPAG), and substantia nigra (SN). Subcortically in the forebrain, we included the bed nucleus of the stria terminalis (BNST) and the nucleus accumbens (NAc). In cortex, we focussed on medial areas 24, 25, 32, 9m, posterior OFC, and frontal operculum (FOP) which, on the basis of their similarities with areas in the monkey brain are most likely to be connected with amygdala ^32^. We also considered the prefrontal areas 46 and 9/46 on the lateral surface because stimulating them both affects amygdala threat-related reactivity ^33^. We used ROIs from the recent parcellation by ^34^ which further subdivides area 24 into a24, p24, and a24pr (the most posterior mid-cingulate region p24pr was not included), area 32 into s32, p32, d32, a32pr, and p32pr, frontal operculum into FOP1-5, and area 9/46 into 9-46d, a9-46v, and p9-46v, and which identifies a pOFC region (for more details, see Methods and **Fig 2C**). For subcortical ROIs, we used established ROIs from published atlases (see Methods) because contrast-based delineation of brainstem nuclei was not available as part of the HCP data.

Fig 2B shows the average functional connectivity from each of the seven amygdala nuclei, merged across hemispheres, to the above-defined 28 cortical, subcortical and brainstem ROIs. While functional connectivity is strongly influenced by the presence of a monosynaptic connection between areas and plastic changes in those pathways, it also reflects multi-synaptic interactions between regions ^35^. Nevertheless, the pattern of functional connectivity observed from the amygdala nuclei exhibited several features reminiscent of animal tracer studies: all amygdala nuclei had strong coupling with areas in ventral, caudal medial frontal cortex and caudal orbitofrontal cortex, including areas 25, pOFC, and s32 as might be expected from non-human primate studies ^19, 36, 37^. Coupling to these regions was strongest for the basal (B, AB) and cortical nuclei (CoN). The same amygdala nuclei had strong coupling with lateral prefrontal regions (46 and 9/46), but the sign was inverted, suggesting negatively correlated BOLD fluctuations. Given the limited connections between the homologue of this region and the amygdala in macaques, it is likely that the negative coupling found between them reflects an indirect interaction mediated by another brain region. In stark contrast, the central (Ce) nucleus had the strongest connectivity to the majority of subcortical and brainstem regions such as NAc or dPAG.

### Behaviour: latent factors capturing mental well-being

Having established and validated the network of connections between amygdala nuclei and our ROIs, we sought a robust characterization of participants’ mental well-being. While the HCP data set is not intended to include patients with clinical diagnoses relating to mental health and is therefore unlikely to include the extremes of the distribution, we reasoned that it might be possible to examine sub-clinical variance in the central range in mental health. We thus selected all behavioural scores available in the HCP data that captured aspects of emotional and psychological well-being, sleep quality, and personality type (see Methods). A total of 33 markers were included which involved measures from the NIH Toolbox ‘Emotion’ (subscales: Psychological well-being; Social relationships; negative affect; stress & self-efficacy), The Pittsburgh Sleep Questionnaire, the Big Five, and the UPenn Emotion Recognition Test. We reasoned that some scores were capturing similar behavioural phenotypes which might have an underlying common cause. To capture such common ‘latent’ factors that produce these mental well-being scores, we performed a factor analysis which resulted in four main factors (see Methods; Fig 3A).

**Figure 3.**
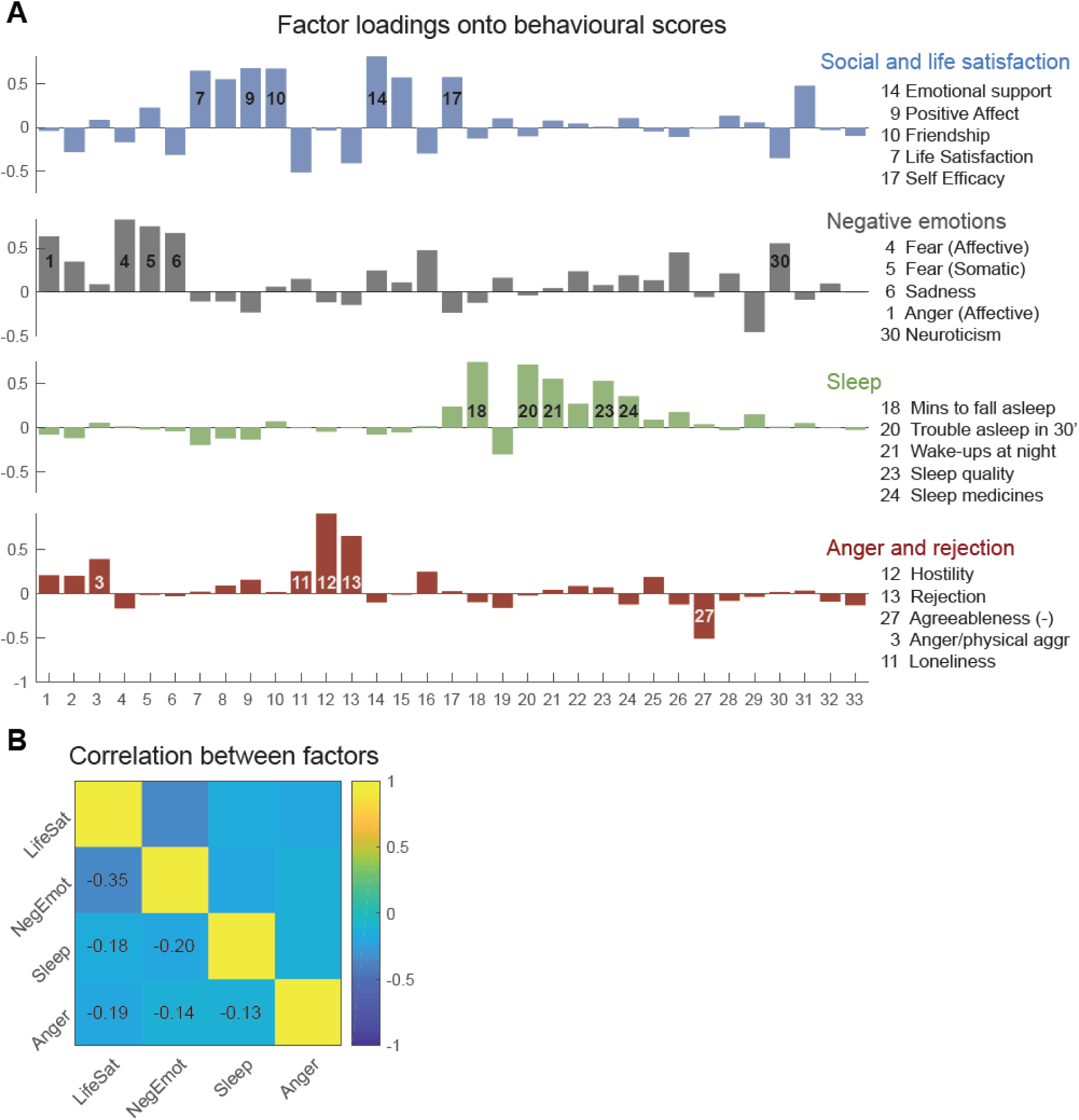
Latent behaviours capture distinct aspects of mental well-being. **A,** A factor analysis conducted based on 33 behavioural scores (**Table 1**) available as part of HCP revealed four factors. The loadings for each factor are shown in different colors, corresponding to the four rows. The highest five contributing behavioural scores are shown in order of their contribution (absolute loading) on the right. This shows that the four factors capture quite distinct aspects of participants’ mental well-being (‘latent behaviours’) which we summarized as ‘Social and life satisfaction’, ‘Negative emotions’, ‘Sleep’ (problems), ‘Anger and rejection’. Importantly, the four factors replicated when the factor analysis was performed on all 1206 HCP participants (see Methods). **B**, Correlations between factors.

The first factor emphasized the impact of social support and general life satisfaction, with a strong negative loading onto loneliness and positive loadings onto emotional support, friendship, life satisfaction and purpose (thus, cutting across the sub-scales of ‘psychological well-being’ and ‘social relationships’ within the NIH Toolbox). The second factor, by contrast, loaded strongly onto negative emotions, including fear, stress and sadness (all within the subscale ‘negative affect’ of the NIH Toolbox). The third factor loaded almost exclusively onto sleep-related markers, assessed as part of the Pittsburgh Sleep Questionnaire. It loaded negatively onto the amount of sleep but positively onto sleep troubles such as bad dreams, wakeups, and lack of sleep quality. Finally, the fourth factor loaded onto anger and physical aggression, hostility, and feelings of being rejected, including negative loadings onto agreeableness (**Fig 3A and Supplementary Fig 2; Table 1**).

**Table 1.** Behavioural markers and their loading onto the four factors

| No | Name | LifeSat | NegEmot | Sleep | Anger/<br>rejection |
| --- | --- | --- | --- | --- | --- |
|  | **NIH Toolbox Emotion Battery (5-point scale)** |  |  |  |  |
| 1 | Anger Affect | -0.04 | 0.63 | -0.08 | 0.21 |
| 2 | Anger Hostility | -0.28 | 0.34 | -0.12 | 0.20 |
| 3 | Anger Aggression | 0.09 | 0.09 | 0.05 | 0.39 |
| 4 | Fear Affect | -0.17 | 0.82 | 0.01 | -0.16 |
| 5 | Feat Somatic | 0.22 | 0.75 | -0.02 | -0.01 |
| 6 | Sadness | -0.31 | 0.67 | -0.04 | -0.03 |
| 7 | Life Satisfaction | 0.64 | -0.10 | -0.20 | 0.02 |
| 8 | Mean Purpose | 0.54 | -0.10 | -0.12 | 0.09 |
| 9 | Positive Affect | 0.67 | -0.23 | -0.13 | 0.16 |
| 10 | Friendship | 0.67 | 0.06 | 0.07 | 0.02 |
| 11 | Loneliness | -0.51 | 0.15 | 0.00 | 0.26 |
| 12 | Perceived Hostility | -0.03 | -0.11 | -0.05 | 0.91 |
| 13 | Perceived Rejection | -0.41 | -0.14 | 0.00 | 0.66 |
| 14 | Emotional Support | 0.81 | 0.24 | -0.08 | -0.10 |
| 15 | Instrumental Support | 0.57 | 0.11 | -0.05 | -0.01 |
| 16 | Perceived Stress | -0.30 | 0.48 | 0.01 | 0.25 |
| 17 | Self-Efficacy | 0.57 | -0.23 | 0.23 | 0.03 |
|  | **Pittsburgh Sleep Questionnaire (scale from 0-9)** |  |  |  |  |
| 18 | minutes to fall asleep (past month) | -0.12 | -0.12 | 0.74 | -0.09 |
| 19 | hours of sleep per night (past month) | 0.10 | 0.16 | -0.30 | -0.16 |
| 20 | sleep trouble: can't go to sleep within 30 minutes | -0.10 | -0.04 | 0.71 | -0.02 |
| 21 | sleep trouble: wake-up in middle of night or early morning | 0.08 | 0.05 | 0.55 | 0.04 |
| 22 | sleep trouble: had bad dreams | 0.04 | 0.23 | 0.27 | 0.09 |
| 23 | overall sleep quality | 0.00 | 0.08 | 0.53 | 0.07 |
| 24 | how often taken sleep medicine | 0.10 | 0.19 | 0.35 | -0.12 |
| 25 | how often trouble staying awake during the day | -0.04 | 0.13 | 0.09 | 0.19 |
| 26 | how often trouble keeping up enthusiasm during the day | -0.11 | 0.45 | 0.17 | -0.12 |
|  | **5-factor model** |  |  |  |  |
| 27 | agreeableness | -0.01 | -0.05 | 0.03 | -0.51 |
| 28 | openness | 0.13 | 0.21 | -0.03 | -0.08 |
| 29 | conscientiousness | 0.05 | -0.45 | 0.15 | -0.03 |
| 30 | neuroticism | -0.35 | 0.55 | 0.01 | 0.02 |
| 31 | extroversion/introversion | 0.47 | -0.09 | 0.05 | 0.03 |
|  | **Penn Emotion**<br>**Recognition Test** |  |  |  |  |
| 32 | number of correct anger<br>identifications | -0.03 | 0.10 | 0.00 | -0.09 |
| 33 | number of correct fear<br>identifications | -0.09 | 0.00 | -0.02 | -0.13 |

We used the loadings from the four factors multiplied onto participants’ original 33 scores to construct latent behaviours capturing these four dimensions of participants’ well-being. We summarized them as ‘social and life satisfaction’, ‘negative emotions’, ‘sleep’ and ‘anger & rejection’.

### Relating latent behaviours capturing mental health to specific amygdala pathways

In the next analysis step, we asked which of the above-defined connections between specific amygdala nuclei and ROIs carried information about mental well-being as captured by the four latent behaviours. For each of the four behaviours, we estimated a large number of regression models using in each case only a subset of connections as predictor variables. This approach has been used, for example, in analyses of human magnetoencephalography data (MEG; ^38^), where recordings across MEG sensors are highly correlated. It is suitable when a large number of correlated regressors precludes simultaneous inclusion in one regression model. Instead of testing each predictor separately, including more than one regressor in each sub-model ensures that variance that is shared among multiple regressors is not attributed to each individual predictor and thus, the unique contribution of each connection can be estimated. More precisely, we estimated *k*=10,000 regression models using a randomly selected subset of 5 out of the total of 196 connections. In each iteration, we recorded which connections were included and we determined the goodness-of-fit using 10-fold cross-validation (CV). In other words, the fit of behaviour achieved using the random subset of five connections was evaluated on 10% of left-out data, and this was repeated ten times, so that predictions were never generated from the same participants that the model was evaluated on (for further details, see Methods). We also ensured that our results were robust to the choice of model size (i.e. the number of connections in each model, here five; see **Supplementary Figure 4**). The contribution of each connection was quantified as the difference in Pearson’s correlation coefficient between predicted and true behaviour (achieved on the out-of-sample data) when the connection was or was not included in the model (*r*Diff; **Fig 4A-C**). An unpredictive connection would have a contribution around zero, meaning its inclusion as a predictor does not boost the performance of the model. By contrast, a predictive connection should improve the correlation between predicted and true behaviour when it is part of the model (positive difference). Overall, the procedure provided a robust quantification of the unique contribution of each connection that was unaffected by existing correlations between predictors (see Methods).

**Figure 4.**
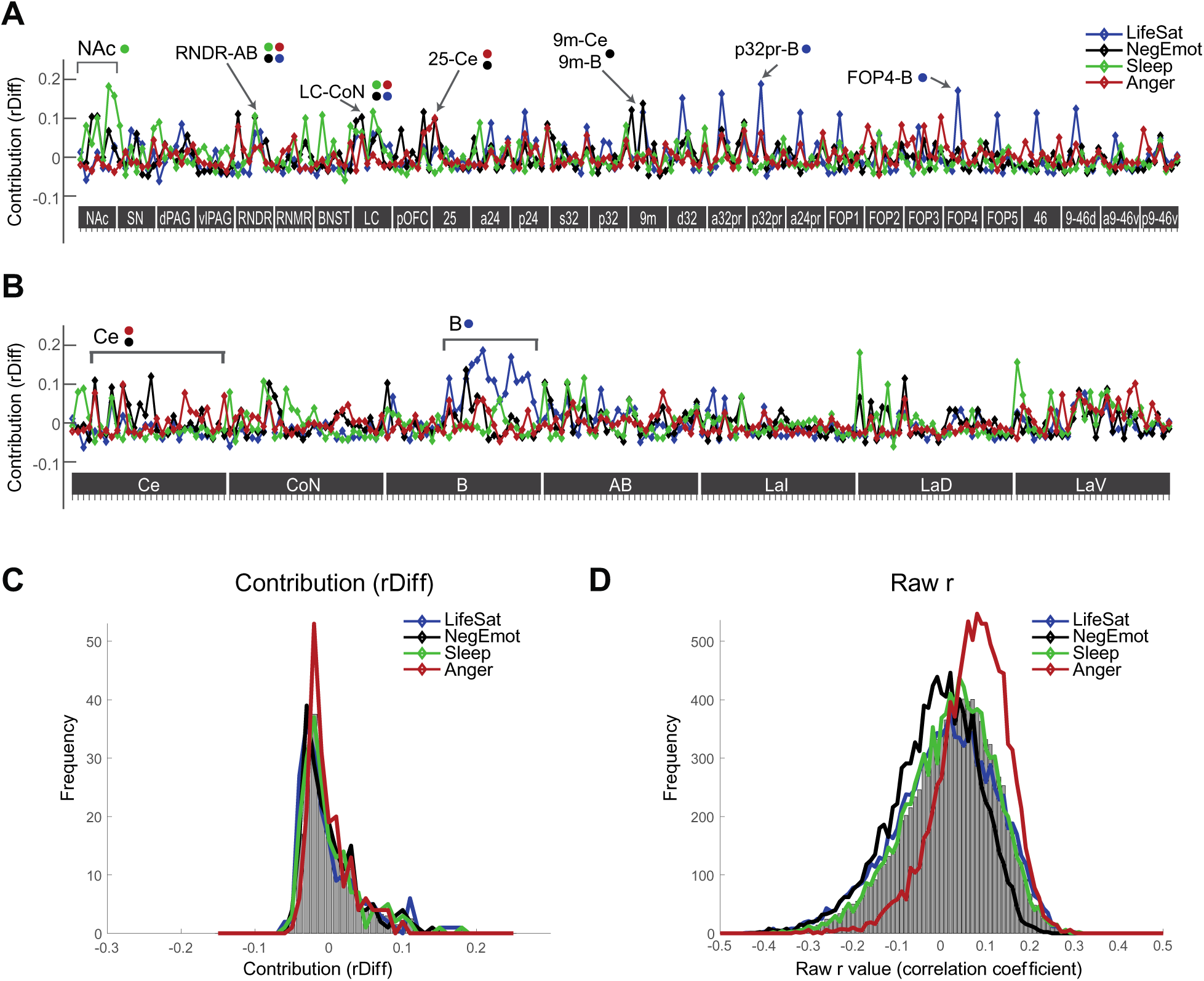
The contribution of specific amygdala connections towards predicting mental well-being. **A,** The contribution *rDiff* of each connection was quantified as the average difference in Pearson’s correlation coefficient *r* between predicted and true latent behaviour when the connection was included in the predictive model versus when it was not included. 10,000 predictive models were run, each including five randomly chosen connections out of the total pool of 196 connections between amygdala nuclei and *a priori* ROIs. All predictions were made using out-of-sample procedures. Visual inspection of *rDiff* values highlights anatomical specificity – e.g. the importance of connections with NAc for predicting sleep, areas p32pr and FOP4 for predicting life satisfaction and some connections predictive of multiple latent behaviours (highlighted with arrows). **B**, Same data as in A sorted by amygdala nuclei instead of ROIs on x. This highlights, for example, the relevance of multiple connections with the basal amygdala nucleus for predicting life satisfaction. **C**, Histogram of contributions *rDiff* across the 196 connections. The majority of connections are unpredictive (around 0). The tail to the right contains predictive connections and shows somewhat stronger predictors for life satisfaction (blue) and sleep (green) than for negative emotions (black) and anger (red). **D**, Histogram of raw Pearson’s correlation coefficients *r* across the 10,000 model iterations.

To provide an intuition for the raw results, we plotted the contribution of all 196 connections for each of the four behaviours, sorted by ROI (Fig 4A) or amygdala nucleus (Fig 4B). Several interesting patterns emerged from visually inspecting these results. First, contributions largely differed between the four behaviours (correlations between pairs of patterns of contributions to the four behaviours, illustrated by the four coloured lines in **Fig 4A-B** were all <.35). For example, the connection between medial dorsal area p32pr and the basal nucleus (p32pr-B) strongly contributed to the prediction of life satisfaction (blue; *r*Diff=.185) but none of the other three behaviours (all *r*Diff<.06), while multiple connections with NAc were relevant for sleep (Nac-LaD: *r*Diff=.179; NAc-LaV=.155) but less for life satisfaction, negative emotions or anger. Second, some ROIs appeared more broadly relevant than others for predicting latent behaviours (more non-zero contributions in Fig 4A): most notably, LC and RN_DR, intriguingly both brainstem nuclei associated with major neurotransmitter pathways, contributed to multiple behaviours via multiple amygdala nuclei. By contrast, some regions, most prominently NAc, already mentioned above, seemed important for a specific behaviour - sleep. Third, examining the contributions sorted by amygdala nuclei highlighted broad differences between amygdala nuclei. For example, the Ce nucleus contributed most to predictions of negative emotions and anger while the basal nucleus was the most critical amygdala nucleus for predicting life satisfaction (Fig 4B). We also examined histograms of, first, the contributions (Fig 4C) and, second, the underlying raw correlation coefficients across the *k*=10,000 regression models (Fig 4D; Supplementary Note 1).

To establish whether contributions of individual connections could be considered meaningful, we performed multiple statistical tests. The first involved comparison against the performance of amygdala connections with randomly chosen other regions. The second was similar but compared our contribution values to those obtained from other subcortical-to-cortical connections selected at random, instead of amygdala connections. The third comparison involved amygdala-to-ROI connections again but this time with the whole amygdala instead of individual nuclei. In a final test, we examined whether binarized behaviours were decodable using single connections.

### Social & Life Satisfaction

For visualization, we sorted connections by the size of their contribution (Fig 5A). We established significance relative to other brain connections in two ways that both corrected for the number of tests. We generated a null distribution based on *n*=1000 different sets of randomly chosen connections, matched in number, and repeated the above model-fitting procedure with a reduced *k*=1000 for computational feasibility. In other words, we ran *k*=1000 iterations of cross-validated regression models each containing five randomly selected connections to predict the original behaviours, and we repeated this for *n*=1000 different sets of 196 randomly drawn connections. In the first test, these randomly drawn connections were always to/from the amygdala (“amy-to-rnd”). In the second test, nuclei of the same size as the amygdala nuclei were defined at random subcortical locations and connections were between these random subcortical seeds and randomly selected cortical regions (“subc-to-rnd”). Thus, in both cases, our control connections were real brain connections and comparable to our original analysis in terms of their signal-to-noise. For each set of random connections, we remembered the contribution achieved by only the top connection. This procedure accounted for multiple comparisons because the same number of predictors as in our original analysis (196) was tested in both of the control cases. The resulting cumulative distribution function was used to establish FWE-corrected p-thresholds (Fig 5A). In the same way, we also generated a distribution based on the contribution *rDiff* of *all* connections in the control hubs, to establish uncorrected p-values (for illustration see **Supplementary Figure 6A**).

**Figure 5.**
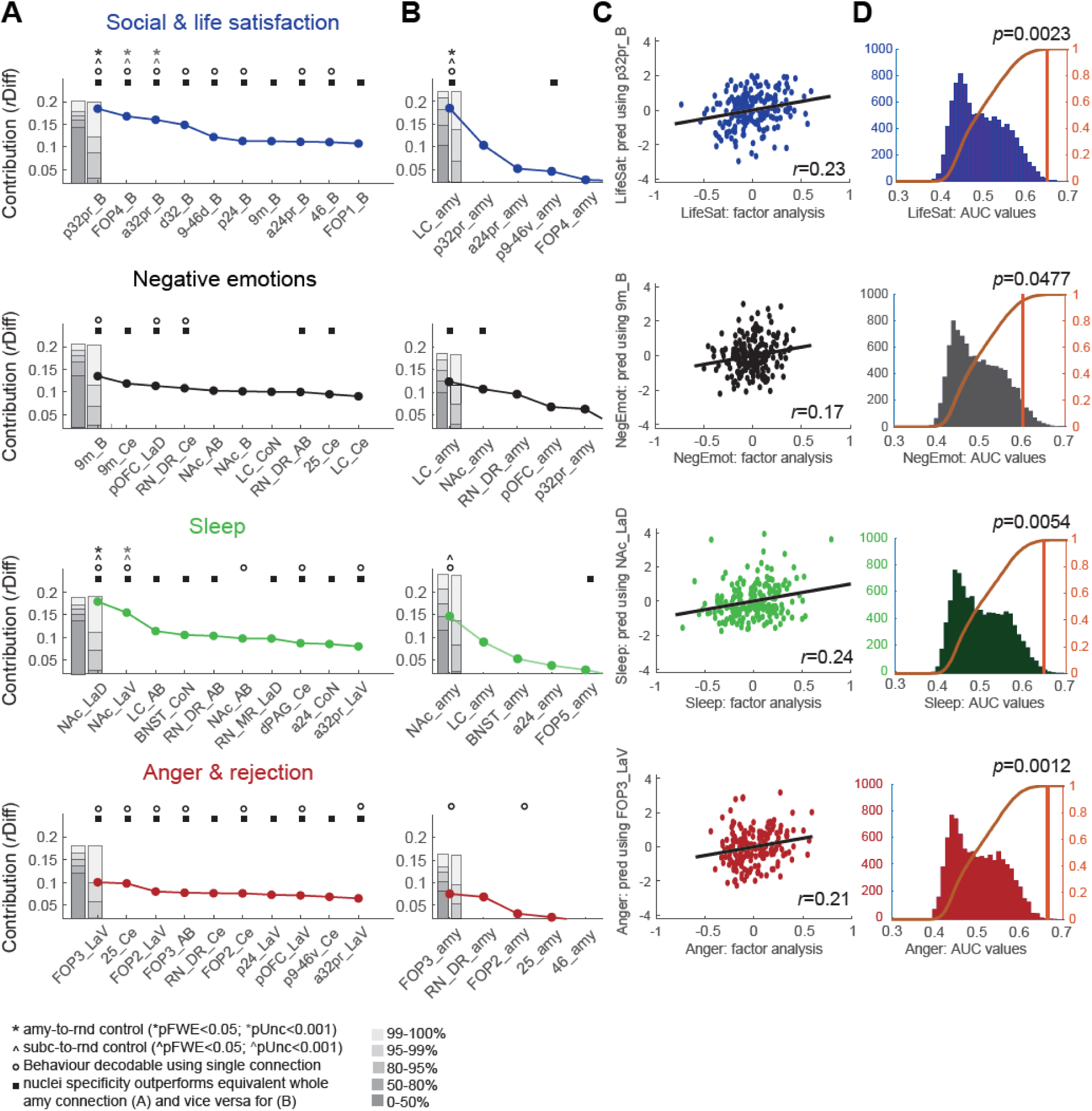
Predictive connections differ across latent behaviours. **A,** Contribution values *rDiff* were sorted for each latent behaviour and the top ten connections are shown in each case. Significance was determined using multiple procedures. First, by considering other amygdala-to-cortical connections (“amy-to-rnd”) and considering the contribution of either only the top connection from the same number of 196 connections (pFWE<0.05: black asterisks *) or all connections (punc<0.001: grey asterisks *). The FWE and uncorrected distributions are illustrated in the two grey bars on the left, respectively. Using the same procedure, a second control was created using random subcortical seeds and their connections to any cortical region (“subc-to-rnd”; pFWE<0.05: black arrow symbols ^; punc<0.001: grey arrow symbols ^). We also tested whether binarized latent behavioural scores (1=top third, 0=bottom third) could be significantly decoded using just a single connection (decodability is denoted by a circle). Finally, we tested whether nuclei-specific connections outperformed the equivalent connection to the whole amygdala (denoted with a square). For predicting social and life satisfaction, connections between the basal nucleus of the amygdala and medial and lateral frontal cortex contributed most. By contrast, subcortical connections and primarily agranular cortical regions were important for predicting negative emotions, and some of these connections were also important for predicting anger. Sleep was predicted by a quite distinct network consisting of connections almost exclusively to subcortical regions. **B**, Contribution values *rDiff* for connections with the whole amygdala, rather than specific nuclei, are sorted according to their *rDiff* contribution and the top five connections are shown. **C**, The true behavioural score obtained from the factor analysis is plotted against the behavioural score predicted, in each case, using only the top connection. This illustrates how a single anatomically specific connection can explain a considerable amount of variance related to a specific marker of mental well-being. **D**, Decodability is demonstrated for the top connection in each case. The AUC value obtained from the top connection is denoted by an orange line; AUC values from shuffled behavioural and connection values are shown in the histogram and were used to generate p-values.

This showed that, for social and life satisfaction, one connection was significant when correcting for 196 comparisons, with two further connections reaching uncorrected significance at p<.001. All three connections were between the basal nucleus of the amygdala and a cortical region: the strongest one with the medial surface of PFC (p32pr with B, p_(FWE,amy-to-rnd)_=.0048; p_(FWE,subc-to-rnd)_=.038), followed by one with frontal operculum (FOP4 with B; p_(unc,amy-to-rnd)_=.0003; p_(unc,subc-to-rnd)_=.0005), and another one with area 32 (a32pr with B, p_(unc,amy-to-rnd)_=.0009; p_(unc,subc-to-rnd)_=.0008; Fig 5A). Their contributions were *rDiff*=.185 for p32pr with B, *rDiff*=.168 for FOP4 with B and *rDiff*=.16 for a32pr with B. Inspection of further connections (Fig 5A) showed that all top connections were with the basal nucleus (B) of the amygdala and cortical regions, predominantly on the medial surface including in addition to the above, d32 with B, p24 with B, 9m with B, a24pr with B, but also some with the lateral surface (9-46d with B and 46 with B). In all cases, a stronger connection between the amygdala and these areas was related to improved life satisfaction (mean regression coefficients all positive: e.g. ß=.226 for p32 and B, ß=.213 for FOP4 and B, Fig 6). Thus, overall, larger coupling values, and thus in many cases weaker negative coupling (Fig 2), between medial and lateral PFC regions and amygdala related to improved life satisfaction. The correlation between the latent behaviour predicted using only the best connection (p32pr with B) and the true latent behaviour is shown for illustration in Fig 5C (*r*=.23).

**Figure 6.**
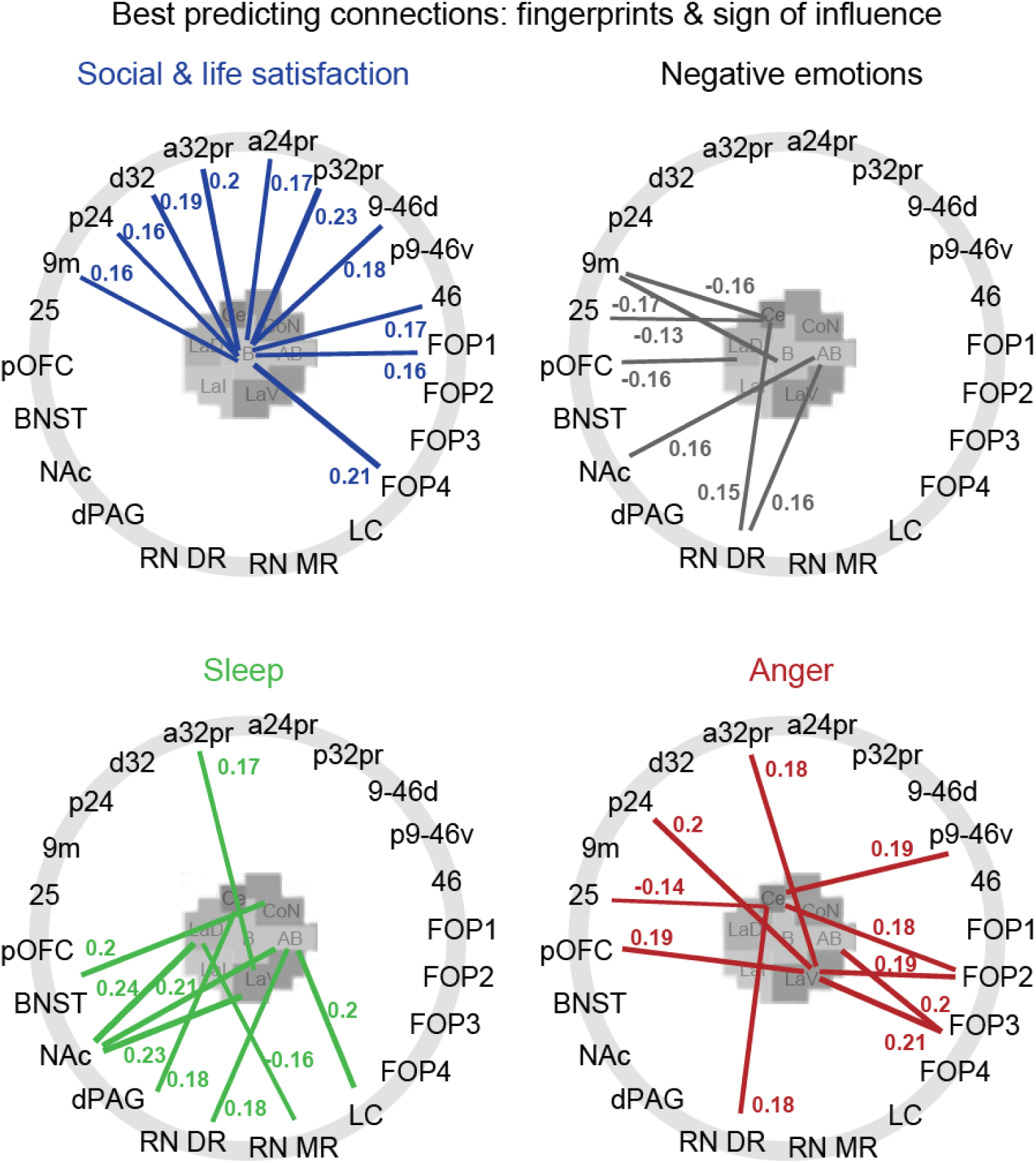
Connectional fingerprints highlight differences between markers of mental well-being. Fingerprints highlight the differences between the connections that significantly contribute to each latent marker of mental well-being. For each connection, the sign and size of influence (regression coefficient) is depicted. For illustration, fingerprints include the top five connections for each behaviour, and any other connections that were significant according to at least one of the four statistical criteria outlined in Figure 5.

In the next step, we tested whether parcellating the amygdala into sub-nuclei increased our specificity for predicting mental well-being. We repeated the above regression procedure for the amygdala as a whole, i.e., using connections of the entire amygdala to the same set of 28 ROIs (**Fig 5B and Supplementary Figure 5**). Again, p-values were obtained using n=1000 repetitions of randomly sampled connections (each with k=1000 models containing five connections). We then used the probability of each whole amygdala connection to set the appropriate alpha level for the corresponding parcellated amygdala connections. In other words, we compared the effect of connections with specific nuclei of the amygdala against the same connections with the amygdala as a whole. If the probability of the parcellated amygdala connection is lower than the threshold set by the whole amygdala, we can infer that the parcellation increased our sensitivity. This showed that, indeed, the nuclei-specific connections to ROIs identified above performed better than would be expected from the connections that reflected connectivity of the same ROI to the entire amygdala. This was true for connections with the basal nucleus compared to the equivalent connection with the whole amygdala for all top connections (denoted with a square in Fig 5A). Only connections with the basal nucleus, but none of the other amygdala nuclei, outperformed whole-amygdala connectivity. Interestingly, the most predictive connection with the whole amygdala, and the only significant one, was with LC (*r*Diff=.185; ß=-.19, *p*_(FWE,amy-to-rnd)_=.0405, *p*_(unc,amy-to-rnd)_=.0014, *p*_(FWE,subc-to-rnd)_=.003, *p*_(unc,subc-to-rnd)_=.0001; Fig 5B and **Supplementary Figure 5**). This connection’s alpha level (i.e. the probability of the same contribution *rDiff* to occur by chance given the control distribution) was smaller than the probability associated with any connection between LC and individual amygdala nuclei (square in Fig 5B). This may seem surprising because in the nuclei-specific analysis, connections with cortical regions seemed most relevant. But in fact, it suggests that for cortical connections with the amygdala, the specificity achieved by subdividing the amygdala into nuclei was crucial for predicting life satisfaction, and all relevant connections were with the basal nucleus. By contrast, the coupling of individual nuclei with LC was not predictive of life satisfaction (compare to Fig 4A), suggesting LC’s interactions with the amygdala are broader and not tied to a specific amygdala nucleus.

In the fourth test, we split life satisfaction scores into thirds and tested whether we could decode whether a participant was in the top or bottom third of participants based on the top connections described above (**Fig 5D and Supplementary Figure 6**). Despite the fact that our investigation focuses on a non-clinical sample that only exhibited limited variation in the behavioural mental health measures, and despite allowing the decoding algorithm to exploit information from only a single connection, prediction accuracies were significant for the top six connections as well as two further connections, including those discussed above (denoted with circles in Fig 5A, see also **Fig 5D and Supplementary Figure 6**; area under the curve (AUC) values, all >.6 and p-values generated from bootstrapping, all p<.05). Thus, using a single anatomically specific connection, in several cases, we were able to decode if someone was more or less likely to be socially connected and more generally satisfied in life.

**Supplementary Figure 3** shows a map of contributions for social & life satisfaction for the entire cortex. The colour in each cortical region reflects the contribution (*rDiff*, z-scored across behaviours and connections) of the functional coupling between the amygdala and that cortical region. In each case it displays the contribution for the amygdala nucleus that was greatest.

### Negative Emotions

The second marker of mental well-being, negative emotions, was subjected to the same statistical tests based on randomly chosen connections with the amygdala or random subcortical seeds, connections with the whole amygdala and decoding of behavioural scores using individual connections. When compared with the distribution of top connections from randomly drawn sets of amygdala (amy-to-rnd) or other subcortical connections (subc-to-rnd), no connection reached significance (FWE-corrected for 196 connections at p<.05 or uncorrected significance at p<.001). Nevertheless, inspection of the identity of the top connections showed that apart from area 9m, the connections that contributed most towards predicting negative emotions were exclusively with subcortical/brainstem regions (RN_DR, NAc, LC) and the most posterior aspects of PFC (area 25, pOFC; Fig 5A), and were therefore clearly distinct from those linked to social and life satisfaction. Interestingly, while connections with all subcortical regions positively related to negative emotions (e.g. ß=.15 and ß=.155 for RN_DR with Ce and RN_DR with AB, ß=.156 and ß=.141 for NAc with AB and BaL, ß=.148 for LC with CoN), stronger positive coupling to medial and orbital PFC regions – which had the strongest influence – was related to reduced chances of experiencing negative emotions (9m with BaL: ß=-.175; 9m with Ce: ß=-.16; pOFC with LaD: ß=-.158, BA25 with Ce: ß=-.129; Fig 6). The correlation between predicted and true behaviour using the best predictor, 9m to B reached *r*=.17 (Fig 5C).

Because the comparisons between amygdala nuclei connectivity and other random selected control connections were not significant after correction for 196 comparisons, the next step of comparing connections of amygdala nuclei with the adjusted alpha level derived from connections of the whole amygdala is perhaps best considered as providing a numerical indication only of whether consideration of individual amygdala nuclei is helpful. Comparison of nuclei-specific connections with those based on whole amygdala demonstrated that some predictions were stronger when they were estimated from sub-nuclei, especially those with cortical regions and RN_DR. For example, predictions achieved from connections with area 9m were better than expected from the corresponding whole-amygdala connection when considering the precise nuclei, Ce and B. The same was true for connections between pOFC with LaD, and area 25 with Ce and RN_DR with Ce or AB. In all these cases, the specific connections’ contributions to a prediction of negative emotion scores were less likely to occur by chance than the adjusted alpha level predicted based on the same ROI’s coupling with the whole amygdala (squares in Fig 5A **and Supplementary Fig 5**). On the other hand, for two subcortical regions, NAc and LC, the adjusted alpha level obtained from the connectivity with the whole amygdala was smaller than the probability of all specific connections with precise nuclei, suggesting that negative emotions can be predicted best from NAc or LC connectivity when considering coupling with the whole amygdala (square in Fig 5B). However, none of the connections with the whole amygdala reached significance (LC: *r*Diff=.123; ß=.15, *p*_(FWE,amy-to-rnd)_=.20, *p*_(FWE,subc-to-rnd)_=.233; NAc: *r*Diff=.107; ß=.14, *p*_(FWE,amy-to-rnd)_=.39, *p*_(FWE,subc-to-rnd)_=.39; **Fig 5B and Supplementary Fig 5**).

Finally, decoding negative emotion scores in the top and bottom third showed that three connections provided significant decoding accuracies, 9m with B, pOFC with LaD, and RN_DR with Ce (circles in Fig 5A, see also **Supplementary Fig 6**; AUCs all >.6, p-values *p*<.05).

### Sleep

Sleep, like social & life satisfaction, was reliably predicted by a subset of amygdala connections when compared with other randomly drawn connections. The connection between NAc and LaV was significant at FWE-corrected levels in both control analyses (*rDiff*=.179, p_(FWE,amy-to-rnd)_=.0039; p_(FWE,subc-to-rnd)_=.014) and another connection with NAc reached uncorrected significance at p<0.001 (NAc with LaD: *rDiff*=.155, p_(unc,amy-to-rnd)_=.0008; p_(unc,subc-to-rnd)_=.0003). Thus, sleep was best predicted by subcortical connections. Inspection of other top connections revealed no cortical predictors in the top eight connections (Fig 5A), but multiple brainstem-amygdala connections that contributed strongly to this behaviour, in stark contrast to both life satisfaction and negative emotions (Fig 6). Both NAc connections related positively to sleep problems. This suggests sleep problems increased with stronger positive coupling between amygdala and NAc (ß=.241, ß=.228; Fig 6). The best connection, NAc to LaV predicted sleep problems with *r*=.24 (Fig 5C).

When considering the coupling between ROIs and the whole amygdala, NAc and LC were the strongest predictors of sleep problems, but neither of them was consistently significant (NAc: *r*Diff=.147, ß=.26, *p*_(FWE,amy-to-rnd)_=.19, *p*_(FWE,subc-to-rnd)_=.008; LC: *r*Diff=.090, ß=.21, *p*_(FWE,amy-to-rnd)_=.80, *p*_(FWE,subc-to-rnd)_=.198). Comparison of nuclei-specific connections with the adjusted alpha level obtained from the corresponding whole-amygdala connection showed for both NAc and RN_DR (as well as all other top connections apart from NAc with B), that the specific connections e.g. between NAc and LaD or LaV, or between RN_DR and AB or LaD were significant given the adjusted alpha level. Thus, they performed better than predicted by chance based on the corresponding whole-amygdala connection. Again, considering sub-regions within amygdala made more specific predictions about mental well-being possible.

Decoding analyses revealed significant decoding for three connections between NAc, with LaV, LaD and AB, but also further connections between dPAG and Ce, and a32pr and LaV (circles in Fig 5A, see also **Supplementary Fig 6**).

### Anger & Rejection

For anger, there was no predictor that performed better, at FWE-corrected levels (corrected for 196 comparisons), than expected from other randomly chosen amygdala or subcortical connections (Fig 5A). Inspection of the top connections revealed a majority of cortical connections, and almost exclusively with the Ce and lateral (LaV) amygdala nuclei. As for negative emotions, cortically, the posterior medial and orbital regions 25 and pOFC were relevant. However, unlike for any other behaviours, the largest number of top contributing connections was with the frontal operculum (FOP2 to LaV and Ce, FOP3 to LaV and AB). Four out of the top six predictors were with frontal operculum. The direction of effects for area 25 to Ce was similar to that seen for negative emotion predictions; increased connectivity between these regions predicted reduced anger (ß=-.14; Fig 6), but stronger frontal opercular connections with the amygdala predicted increased problems with anger (ß=.207, ß=.188, ß=.202, ß=.183). The correlation between predicted and true behaviour based on the best predictor, FOP3 to LaV, was *r*=.21 (Fig 5C).

The best connections between ROIs and the whole amygdala matched those identified above in terms of their ROI targets for anger (FOP3: *rDiff*=.076, ß=.17, *p*_(FWE,amy-to-rnd)_=.56, *p*_(FWE,subc-to-rnd)_=.33; RN_DR: *r*Diff=.067, ß=.18, *p*_(FWE,amy-to-rnd)_=.66, *p*_(FWE,subc-to-rnd)_=.42). Interestingly, in all cases, e.g. for connections between area 25 with Ce, FOP2 with LaV or Ce, and p24 and pOFC with LaV, the probability of nuclei-specific connections was smaller than predicted from the adjusted alpha level derived from the corresponding whole-amygdala connection, showing that the specificity provided by the nuclei improved the prediction (squares in **Fig 5A and Supplementary Fig 5**).

Decoding binarized anger scores showed that multiple of our top connections allowed above-chance predictions, most prominently those with FOP regions, namely FOP3 with LaV and AB, and FOP2 with LaV and Ce, but also area 25 with Ce, pOFC with LaV and a32pr with LaV (all AUC>.6 and p<.05).

## Discussion

The need to better describe the biological underpinnings of psychological illness and dimensional variation linked to psychological illness has long been recognized (for a recent perspective, see ^23^). Here, we used resting-state fMRI, a common *in vivo* tool for estimating human brain connectivity, but applied a fundamentally different rationale and approach to the analyses of both neural and behavioural data. Turning first to behavioural data analysis, rather than stratifying a disease such as depression into several biologically meaningful sub-groups ^39^ or classifying people into categories (e.g. patient vs control), we aimed to define biologically meaningful latent behaviours that capture central aspects of mental health that exhibit variation even in the sub-clinical range. In the neural analysis we were able to predict these latent behaviours using a small number of anatomically motivated brain connections. All of this was done using out-of-sample methods, ensuring robustness and internal replicability.

We identified four latent behaviours which we believe capture distinct aspects of people’s mental health: social/general life satisfaction, negative emotions, sleep problems, and problems with anger/rejection. Rather than using a summary measure, such as e.g. the total depression score, we reasoned that because specific brain connections carry specific combinations of input and output, mappings of behaviour onto precise brain connections are more likely achieved for functionally meaningful behavioural units ^23, 40^. We obtained latent behavioural markers using a factor analysis ^41^. Alternatively, computational modelling approaches are sometimes used to identify precise measures of behaviour. ^42, 42–44^ In order to link the latent behavioural markers to precise brain connections, we focussed on the amygdala. First, we demonstrated that it was possible to identify *in vivo* seven component amygdala subregions that corresponded to amygdala nuclei. They reliably varied in their connectivity in comparison to one another, but they were topological arranged in a similar manner in both hemispheres. Second, we demonstrated that patterns of functional connectivity – correlations in the BOLD signals – between each amygdala nucleus and 28 cortical, forebrain subcortical, and brainstem regions were approximately as predicted from anatomical tracer studies. We were then able to proceed to the final stage of the study and show that variation in functional connectivity between specific amygdala nuclei and these other regions were predictive of variation in the four latent behaviours.

Three aspects of our data underlined the importance of the functional connectivity of specific amygdala nuclei. First, several of the best predictive connections associated with each latent behaviour explained enough variance to allow prediction of whether someone was in the top or bottom third of the given behaviour. The resulting decoding accuracies achieved using a single connection (**Fig 5A,D** and **Supplementary Fig 6**) were not too dissimilar from accuracies reported when predictions were based on large networks ^1–5^ and we would expect them to be even larger in a clinical population that includes the extremes of the behavioural distribution. In the context of neuroimaging, our sample size of 200 participants can be considered fairly large ^23^. We therefore believe the reported effect sizes are considerable and meaningful. Despite the importance of large network approaches, an advantage of the current approach is that it provides specific regions and connections as targets for therapeutic intervention involving a range of approaches such as pharmacological, neurostimulation, neurofeedback, or cognitive interventions. Second, variations in six or more of the connections were associated with significantly better predictions of the latent behaviours than was possible when just the connectivity of the amygdala as a whole was considered. Finally, in a third test, we established that the connection’s contributions were significant even when the null distribution that they were compared against was from the same amygdala nuclei but to a random set of 28 brain regions, or from same-size random subcortical nuclei to a random set of 28 brain regions. Indeed, for two of our latent behaviours, two to three connections between specific amygdala nuclei and other brain regions were significant predictors of the extent to which the behaviour was present, over and above what would be achieved using randomly chosen connections. Importantly, predictive connections largely differed between the four latent measures of mental well-being (see fingerprints in Fig 6) and only few connections were shared.

We had a strong anatomical prior not only on the importance of the amygdala but the importance of the amygdala’s interactions with specific cortical, forebrain subcortical, midbrain, and brainstem regions thanks to the large body of studies in animal models that has examined these circuits ^10–13, 20^. As a result of careful fMRI data preprocessing we were able to examine activity not just within medial temporal lobe but even in specific brainstem regions and relate the coupling patterns to variation in our indices of mental health. We believe there are other prime anatomical hubs such as ventromedial and subgenual frontal areas that would be worth investigating with a similar approach. It is unlikely that a single region or network is sufficient to fully predict all aspects of someone’s mental well-being ^7, 15, 20, 39, 45, 46^. Nevertheless, we believe it is important to recognize that individual and identifiable connections may have particular importance. This is a view taken more commonly when considering targeted interventions in mental health, such as for example using invasive deep brain stimulation (DBS) which has in some cases led to remarkable improvements in mood ^45, 47^, but may work particularly well when the right connections between subcortical and cortical regions are targeted ^48^. Similarly, other non-invasive stimulation approaches such as repetitive transcranial magnetic stimulation (rTMS) are more likely to be successful when targeting specific circuits (e.g. subcallosal connectivity; ^49^). Such interventions could become more feasible with advances in non-invasive ultrasound methods ^50–52^. So, while our findings are not of immediate clinical relevance, they suggest interventions targeted at particular nuclei might benefit someone predominantly suffering from sleep problems while targeting others might benefit someone who experiences strong negative emotions. We note one potential limitation, namely that we relied on a large volume of data – approximately one hour of resting-state scans in each participant – from highly optimized pulse sequences, which may not be available regularly in patients.

Our parcellation of the amygdala into seven nuclei strikingly resembled previous amygdala investigations but which were possible only *post mortem* ^29–31^. Saygin et al., for instance, scanned at a resolution of 100-150um at 7T and identified nine nuclei which resembled in their size, position and transitions patterns the seven nuclei identified here. Previous parcellations based on *in vivo* data have identified fewer subdivisions ^24–27^ but the borders identified in those studies still resembled a subset of the borders we identified here thereby underlining consistency in results. The finer grained parcellation we obtained reflected improved image quality and preprocessing pipelines that better controlled for physiological noise. We show that detailed amygdala parcellation is important for achieving the behavioural prediction accuracies reported here (Fig 5). This is unsurprising given known anatomical and functional differences between amygdala nuclei ^19^.

The amygdala networks identified for the different latent behaviours seem plausible in the context of previous work. For example, social and life satisfaction highlighted connections between the amygdala and regions primarily located in medial and lateral frontal cortex, more precisely areas p32pr, a32pr and d32 as well as areas FOP4 and 9-46d ^34^, with less pronounced negative coupling between these areas and the basal amygdala nucleus predicting improved life satisfaction. These areas in or close to the dorsal anterior cingulate cortex (dACC) as well as frontal opercular/insula regions have been linked to aspects of behavioural change and adaption ^53, 54^, abilities compromised in anxiety ^55^, and are important for arbitrating between exploration and exploitation ^56, 57^, a process changed in depression ^58^. Even though there is probably little direct coupling between dlPFC and amygdala, dlPFC is a stimulation target in depression, and alters amygdala threat responses ^33^. It thus seems unsurprising that connections between these medial and lateral frontal regions and the amygdala might contribute to overall social and life satisfaction.

It is worth noting that, because rs-fMRI was used as a proxy for anatomical connectivity here, the patterns in activity coupling we identify do not necessarily correspond to monosynaptic connections. While monosynaptic connections might dominate in Figure 2B which illustrates, the strongest activity correlations of the amygdala nuclei, the relations we identified between activity coupling and mental health indices (Figures 4-6) may rely on a multi-component connection pathway or may involve connections between two amygdala nuclei.

The associations between negative emotions, our second latent behaviour, and amygdala connectivity can also be understood in the context of the functions of these areas even if, once again, some of the critical pathways may be indirect. Amygdala connections with areas 9m, pOFC, 25 and subcortical structures (LC, RN_DR, NAc) seem plausible. Weaker coupling between area 25 and the central nucleus, between pOFC and the adjacent LaD nucleus, and between 9m and Ce and B nuclei are related to more pronounced negative emotions. Although pOFC has received little attention, bipolar patients demonstrate reduced grey matter in pOFC ^59^ and both pOFC and amygdala have been linked to the most basic aspects of stimulus-reward association learning ^60^. Area 25 has been linked to autonomic and affective regulation and, just like the amygdala, exhibits abnormal metabolism in depressed patients ^15, 20^. Stimulation of this region or its interconnections may reduce depression ^45, 48^. The fourth latent measure of mental well-being we identified, anger and rejection, was also linked to areas 25 and pOFC, in addition to other frontal opercular regions that have recently been linked to the balancing of the most recent outcomes with the wider, more long-term experience of reward ^61, 62^.

In contrast to cortical regions, stronger rather than weaker subcortical connectivity with amygdala nuclei predicted negative emotions. This suggests that diminished cortical-amygdala interaction is accompanied with increased amygdala interaction with subcortical areas linked to the origins of widely branching neuromodulatory systems such as serotonin and noradrenaline (RN_DR, LC) and key targets of other systems such as dopamine (NAc). Noradrenaline mediates stress and stress-related responses and stress-induced dysregulation of the NA system may contribute to the pathogenesis of depression ^63^. Increasing NA can also be effective as an antidepressant treatment. LC occupied a somewhat unique position because it was the only region which was somewhat predictive of three out of four latent behaviours when considering coupling with the whole amygdala (**Fig 5B and Supplementary Fig 5**). This suggests that a more global coupling pattern between LC and amygdala may help regulate mood in a way that impacts multiple of our latent measures of well-being. Indeed, LC-amygdala coupling has been linked to the retrieval of emotional memories ^64^. Taken together, LC-amygdala connections seem central for mediating problems related to negative emotions that impact mental health.

The third latent measure of mental well-being captured sleep problems and was linked to a distinctly different connectional fingerprint (Fig 6). Unlike the other three behaviours, it comprised only subcortical connections between lateral amygdala nuclei and NAc. The NAc is an important projection target of VTA dopamine neurons, and dysfunction of the striatum has been associated with sleep disturbances, with neurons in NAc core particularly important for controlling slow-wave sleep ^65, 66^.

In summary, our work suggests that strong anatomical priors derived from animal studies, in combination with neuroimaging data of sufficient anatomical detail, make it possible to forge links between dimensions of mental health and specific neural circuits. Crucially this also depends on the identification of mental health behaviour clusters which, even if in the subclinical range, are naturally emerging functional groupings that are more likely to map onto the brain’s functional organization.

## Materials & Methods

### Participants

Data and ethics were provided by the Human Connectome Project (HCP), WU-Minn Consortium (Principal Investigators: David Van Essen and Kamil Ugurbil; 1U54MH091657) funded by the 16 NIH Institutes and Centers that support the NIH Blueprint for Neuroscience Research; and by the McDonnell Center for Systems Neuroscience at Washington University. Two hundred HCP subjects (n=200; mean age 29 ± .26; age range 22-36; 108 females, 92 males) were pseudo-randomly chosen from the full HCP data set (https://www.humanconnectome.org/). Working on a subset of all 1206 HCP participants was necessary because one key aspect of the pre-processing was to correct rs-fMRI data for physiological noise, which particularly affects the key regions of this study such as the amygdala and brainstem. However, the quality of acquired physiological variables varies substantially across HCP participants. We therefore inspected the variance in physiological recordings of those participants’ in whom physiological measures had been acquired both visually and by plotting summary measures such as the total variance over time and only considered participants with sufficient signal in both cardiac and respiratory measurements. Participants were further selected to achieve a spread in their mental well-being scores. Specifically, we tried to achieve high variance in the total DSM score (ASR_Totp_T) which was not used in any further analyses (resulting mean total DSM score: 47.94, variance: 103.83; mean of all 1206 HCP participants: 47.41; variance: 80.61).

### Data and minimal pre-processing

Four resting state runs were acquired on a Siemens Skyra 3T scanner using custom pulse sequences (for details see ^67–69^). In brief, resting-state runs lasted 14.4 minutes, had a TR of 720ms, TE of 33ms, isotropic resolution of 2mm, 72 slices, and a multiband factor of 8 resulting in 1200 timepoints. Two runs were acquired using right-left phase encoding and two using left-right phase-encoding. Spin-echo images and T1-weighted images were acquired for distortion correction and registration (for more details see ^70^). We used all four runs of each subject and downloaded the minimally pre-processed HCP data which is described in detail in ^28^. In brief, these data are distortion-corrected, temporally filtered, projected on to a surface reconstruction obtained from the T1-weighted image while maintaining subcortical voxels (cifti format), and minimally smoothed. Registration across participants was achieved using multi-modal areal-feature-based surface registration (MSMall) ^34^.

### Additional pre-processing

Because noise caused by physiological artefacts (e.g. breathing, pulse) is particularly pronounced in brainstem and temporal lobe structures, all key areas for this study, we performed corrections for physiological noise in the data. Removal of artefacts caused by physiological signals is not currently incorporated in standard HCP pipelines. We used the PNM toolbox (https://fsl.fmrib.ox.ac.uk/fsl/fslwiki/PNM; ^71^) to generate physiological regressors (a total of 33 regressors comprised of: cosine and sine of basic cardiac and respiratory regressors modelled with an order of 4, and thus 16 regressors; multiplicative cardiac and respiratory terms cos(c+r), sin(c+r), cos(c-r), sin(c-r), each modelled using an order of two, and thus again 16 regressors; plus respiration volume per time (RVT)^71^). In addition to physiological regressors, we constructed 24 motion regressors from the six motion regressors provided (in the HCP data release, these are stored in Movement_Regressors.txt) (e.g., ^70^): the six original regressors, their derivatives, and the square of the resulting twelve regressors. We also used independent component analysis (ICA)-denoising as provided with the ‘fixextended’ HCP dataset (melodix_mix and Noise.txt). The motion, physiological and ICA noise regressors were normalized, high-pass filtered and detrended to mimic the pre-processing performed on the data. Then, motion and physiological confounds were aggressively regressed out of the data and ICA components (thus entirely removing any variance explained by physiological or motion parameters), and the noise ICA components were subsequently removed from the data using a soft regression (thus removing only the variance unique to the ICA noise components).

The data were demeaned, the variance of the noise in the data normalized (as in ^34^) and the four runs of each participant were concatenated. Additional smoothing was applied to the surface only (sigma=5mm; no additional smoothing was applied to subcortical structures, including the amygdala). This yielded the fully pre-processed data for each of the 200 participants which contained a total of 4800 time points from the combined 1200 time points of the four resting-state runs.

### Group dense connectome

A group average timeseries was generated from the 200 individual data sets using the algorithm ‘MIGP’ ^72^. MIGP is a computationally tractable method to approximate the group average time series using group-level PCA. The two parameters specifying (a) the number of data-points kept on-line during the iterative computation of the average and (b) the cut-off describing the number of principal components kept at the end were both set to 4800, corresponding to the number of data points in each individual’s file. A dense connectome was created from the average time series using the function cifti-correlation (using Fisher’s z). Ringing artefacts were corrected using Wishart RollOff ^34^.

### Clustering

The full dense connectome was restricted to contain the connectivity of voxel’s in both amygdalae to the rest of the brain (647 voxels × 91282 brain-ordinates). Connectivity values were transformed into absolute values (i.e., unsigned ‘strength’ of correlation) to enable both positive and negative coupling patterns to inform the clustering solution (FSLnets ignores negative values in its hierarchical clustering routine). A similarity matrix was computed based on this absolute connectivity using Pearson’s correlation coefficient (FSLnets function nets_netmats, part of FSLnets: https://fsl.fmrib.ox.ac.uk/fsl/fslwiki/FSLNets). In other words, the similarity matrix captured, for any pair of amygdala voxels, the similarity of their connectivity profile to the rest of the brain. The similarity matrix was fed into a hierarchical clustering algorithm (function nets_hierarchy.m part of FSLnets). We thus obtained a clustering of the amygdalae based on the similarity of different amygdala voxel’s connectivity to the rest of the brain.

To evaluate the number of clusters, or in other words, the appropriate depth of the hierarchical clustering tree, we aimed for a good balance between simplicity and detail, as well as anatomical plausibility. One simple heuristic to assess anatomical plausibility was to prefer solutions with corresponding clusters across left and right hemispheres. Another focus was on detail: for instance, it has been suggested that the two largest amygdala nuclei, basal and lateral nuclei, can be further split into several subdivisions ^31^. We were also keen to identify the rather small central nucleus in both hemispheres, given its importance for connecting the amygdala to brainstem regions. The central nucleus split off at depth 10 and 12 of the hierarchical clustering algorithm in the left and right hemisphere respectively, at which point both hemispheres contained 7 clusters (AB and CoN were still connected across hemispheres, so there were five uniquely left clusters, five uniquely right clusters and two clusters that contained both hemispheres, and thus depth 12). This clustering solution was also symmetrical across hemispheres. At the next depths from 13-15, AB split between L and R hemispheres and the ventral part of the lateral nucleus split into two halves first in the right hemisphere and then in the left hemisphere. This was more detail than we would have anticipated or required for interpretation of further analyses. Throughout the results, we therefore focussed on the depth 12 cluster solution, which when merging corresponding clusters in both hemispheres yielded seven final clusters (**Figs 1B and 2A**). Other clustering depths are shown in **Supplementary Fig 1**.

### Naming of clusters

The labelling of clusters was largely based on the Atlas of the Human Brain by ^31^ and a post-mortem parcellation at 7T with 100-150um resolution ^29^. The most dorsal, posterior and lateral nucleus (dark blue in **Figs 1B and 2A**) which was also the smallest in size (62 voxels across both hemispheres) perfectly matched in its size and position the central amygdaloid nucleus and was therefore labelled **Ce**. Judging from its position and size, it contained both medial and lateral divisions of the central nucleus’ ^31^. However, it is less clear whether it also contained the medial amygdaloid nucleus. The medial amygdaloid nucleus might have been part of this ‘Ce’ cluster or the adjacent cluster (middle blue in **Figs 1B and 2A**) which was positioned in a dorsal, posterior and medial location where the cortical amygdaloid nuclei are located (e.g. PCo=posterior cortical; ACoV and ACoD = anterior cortical, ventral & dorsal parts ^31^; sometimes referred to as CAT = cortico-amygdaloid transition area e.g., ^29^). We therefore labelled this adjacent cluster **CoN**, as an agglomeration of the cortical nuclei of the amygdala. It contained altogether 133 voxels across left and right hemispheres, and possibly comprised cortical nuclei as well as the medial nucleus. Ventral and anterior to the Ce and CoN nuclei, in a medial position within the amygdala (light blue in **Figs 1B and 2A**), we identified a portion of the basal amygdala which very likely contained Mai et al.’s ventral and dorsal basomedial (BMVM and BMDM), and probably also its basolateral paralaminar and intermediate subdivision (BLPL and BLI), and thus the majority of basomedial and basolateral aspects of the basal nucleus. We therefore refer to it simply as the basal nucleus **B**. It contained 74 voxels and was adjacent to a slightly more medial subdivision of the basal nucleus which we refer to as auxiliary basal (**AB**, green in **Figs 1B and 2A**) and which contained 104 voxels. This cluster AB, based on its size and location, would have contained the ventromedial part of the basolateral nucleus in ^31^ (BLVM) and closely corresponded to what Saygin and colleagues ^29^ describe as AB as well. The remaining three clusters made up the lateral nucleus of the amygdala, namely its dorsal, intermediate and ventral portion (**LaD**, **LaI**, **LaV**, respectively, in red, yellow and dark red in **Figs 1B and 2A**). These clusters contained 84, 86 and 104 voxels, respectively.

### ROI selection

We had a number of a priori regions of interest which were informed by prior work, including anatomical work using tracers in macaque monkeys as well as work in humans with mental health disorders. All our ROIs are illustrated in Fig 2C and will be motivated one by one.

We included aspects of dorsolateral prefrontal cortex (dlPFC) despite it not having strong monosynaptic connections with the amygdala in monkeys, because of work involving neurostimulation to dlPFC, most commonly repetitive transcranial magnetic stimulation (rTMS), which has been shown to alleviate symptoms of mental health disorders, particularly depression ^73, 74^ and because it has been implicated in the regulation of amygdala responses to threat ^33, 75^. The location of stimulation over dlPFC can be variable across studies but is most common over areas 9/46 and 46 and particularly effective when strong connectivity with dlPFC and area 25 is observed ^76^. We therefore included all sub-clusters of areas 46 and 9/46 reported in HCP’s multi-model parcellation version 1.0 ^34^ which included 46 (316 vertices), 9-46d (379 vertices), a9-46v (147 vertices), and p9-46v (214 vertices).

On the medial and orbital surface, amygdala connectivity gradually changes along an anterior-posterior axis, with strongest connectivity posteriorly closest to the corpus callosum ^19, 77^. This also mimics the transition between agranular and dysgranular/granular cortex, and unimodal to transmodal connectivity ^78, 79^. We included all agranular regions in the medial and orbital prefrontal cortex; all of the likely homologues of these areas have strong monosynaptic connectivity with the amygdala in monkeys. This included areas 32, 25, 24 and the most posterior part of OFC. We also included granular area 9m, adjacent do area 32 in medial frontal cortex, and frontal operculum, which has also been highlighted in tracer studies for its connections with the amygdala. As above, we took the parcels obtained from HCP’s multi-model parcellation version 1.0 ^34^ which are labelled areas 25 (54 vertices), a24 (89 vertices), p24 (66 vertices), a24pr (75 vertices), s32 (55 vertices), p32 (122 vertices), d32 (147 vertices), a32pr (163 vertices), p32pr (190 vertices), 9m (408 vertices) and pOFC (83 vertices). Frontal operculum contains FOP1-FOP5 (with 61, 101, 83, 240 and 193 voxels, respectively). Apart from their strong connectivity with the amygdala many of these regions have indeed been implicated in mood disorders and social cognition. For example, PET work shows abnormal metabolism in subgenual PFC, including area 25, ventral 24 and possibly 32 ^15^, deep brain stimulation in sub-genual regions or their adjacent fibre passages can alleviate symptoms of depression ^45, 80, 81^ and negative biases in decision-making can be induced by stimulating pregenual cortex (rostral area 24 and dorsal area 32; ^82^). In addition, several studies have also highlighted the importance of peri-/pregenual cortex in social cognition ^83, 84^. In summary, we included four cortical regions on the lateral, 10 cortical regions on the medial, one cortical region on the orbital surface, and five frontal opercular regions, and thus a total of 20 cortical ROIs.

Subcortically, our major focus was on the key nuclei associated with different neurotransmitter systems because of their importance for mental well-being. We included the substantia nigra (SN), which contains the majority of dopaminergic neurons and the nucleus accumbens (NAc), an area receiving strong dopaminergic innervation ^85^. DA has been implicated in mental health disorders; for example, Parkinson’s disease, which is characterized by a loss of DA neurons in SN, leads to depression in a large percentage of patients (∼35%; ^86^). But DA also plays a key role in reward-learning and sleep regulation. Striatal dysfunction has, for instance, been associated with sleep disturbances and a subset of NAc core neurons was found to regulate slow-wave sleep ^65, 66^. The SN mask was taken from the NITRC Atlas of the basal ganglia ^87^ and contained 134 voxels. The NAc was taken from the Harvard Subcortical Atlas and contained 188 voxels.

The bed nucleus of the stria terminalis (BNST) was included because of its role in mediating the long-term effects of anxiety and responses to stress ^88^. It is also sometimes considered part of the extended amygdala. The BNST mask was obtained from ^89^.

Two regions with opposing functionality within the periaqueductal grey (PAG) were included because of their importance in regulating autonomic arousal: ventrolateral PAG (vlPAG) which mediates rest- and digest-related behaviour and dorsal PAG (dPAG) which mediates fight and flight responses. The masks for these regions were taken from ^90^. The dPAG was the summation of Faull et al.’s dorsomedial (dm) and dorsolateral (dl) aspects of PAG; vlPAG contained 43 voxels and dPAG 45 voxels.

The role of serotonin and of selective serotonin reuptake inhibitors (SSRI) in the pathology and treatment of mental health disorders is well known. The raphe nuclei are the most important source of serotonin in the brain. Masks for dorsal and median raphe nuclei were taken from the Harvard Ascending Arousal Network Atlas ^91^. The dorsal raphe nucleus (RN_DR) contained 23 voxels, and the median raphe nucleus (RN_MR) contained 8 voxels. Finally, locus coeruleus (LC), the main site of noradrenaline production was defined based on ^92^ and contained 20 voxels.

Probabilistic masks were binarized first, including all voxels with probability >.25, in other words, voxels that had a larger than 25% chance of being within the given region (NAc, SN). Binary files and all masks we received in binary format (BNST, PAG, LC, RN) were subsampled to 2mm, and binarized again using any voxels >.25 in subsampled space. The exceptions were NAc where thresholding at .25 would have yielded an unusually large ROI, so a threshold of >.75 was applied in the second step; for the raphe nuclei, thresholds were adjusted manually to maximise anatomical plausibility (>.6 and >.72 for dorsal and median, respectively).

Thus, we included a total of eight subcortical and brainstem regions, which are shown in Fig 2C.

### Selection of behavioural scores

Instead of using psychiatric scores (e.g. the total depression score), our goal was to define underlying variation in emotional and social wellbeing in the normal range but, in particular, those aspects of emotional and social wellbeing that might be affected in anxious or depressed individuals. We went through all restricted and unrestricted behavioural markers acquired as part of HCP and selected those that related to mental well-being. We included 33 behaviours composed of

1. Measures from the NIH Toolbox Emotion Battery (www.nihtoolbox.org) ^93, 94^(total: 17); each item was administered on a 5-point scale with options ranging from “not at all” to “very much”. In all cases a-d below, scores <40 are considered low and scores >60 are considered high.

a. a. Six measures from the Negative Affect toolbox (Anger Affect: obtained using computer-adaptive testing (CAT), Anger Hostility: obtained from a questionnaire with 5 items, Anger Aggression: also 5 items, Fear Affect: CAT, Fear Somatic: 6 items, Sadness: CAT);
b. b. Three measures from the Psychological Well-Being toolbox (Life Satisfaction, Mean Purpose, Positive Affect) – all obtained using CAT
c. c. Six measures from the Social Relationships toolbox (Friendship, Loneliness, Perceived Hostility, Perceived Rejection, Emotional Support, Instrumental Support); loneliness obtained from a questionnaire containing 5 items, all others from questionnaires containing 8 items.
d. d. Two measures from the Stress and Self Efficacy toolbox (Perceived Stress: 10 items, Self-Efficacy: CAT)
2. Measures from the Pittsburgh Sleep Questionnaire ^95^ (total 9) composed of minutes to fall asleep (past month); hours of sleep per night (past month); sleep trouble: can’t go to sleep within 30 minutes; sleep trouble: wake-up in middle of night or early morning; sleep trouble: had bad dreams; overall sleep quality; how often taken sleep medicine; how often trouble staying awake during the day; how often trouble keeping up enthusiasm during the day. All of these were rated on a scale from 0-9.
3. Measures from the five-factor model ^96^ a) neuroticism; b) extroversion/introversion; c) agreeableness; d) openness; and e) conscientiousness ^97^. HCP data collection administered the 60-item version of the Costa and McRae Neuroticism/Extroversion/Openness Five Factor Inventory (NEO-FFI), which has good reliability and validity ^97^. This measure was obtained as part of the Penn Computerized Cognitive Battery ^98^.
4. Measures from the Penn Emotion Recognition Test, again obtained as part of the Penn Emotion Recognition Test. During this test, participants are presented with 40 faces and need to identify the emotion of the face from the five options happy, sad, angry, scared and no feeling. There are eight faces in each category. We included a) the number of Correct Anger Identifications (ER40ANG) ranging from 0-8 and b) the number of Correct Fear Identifications (ER40FEAR) ranging from 0-8.

### Factor analysis and creation of latent behaviours

We conducted a factor analysis on these 33 behavioural markers (z-scored) using Matlab’s function ‘factoran’, with a ‘promax’ rotation. A Scree test ^99^ based on an initial sample of 100 participants suggested four factors (nFactors package in R with function nScree ^100^), all of which seemed interpretable upon inspection of their weights. We therefore fixed the number of factors to four. Importantly, the same four factors replicated in our full dataset of 200 participants and inspection of a potential fifth factor showed lack of interpretability and would have introduced a high correlation between two of the factors (r=.5; compared to highest correlation in our set of four: .35). Moreover, and most importantly, our four factors also replicated on the full set of 1206 HCP participants: the correlation between the factor weights for a factor analysis based on 200 versus 1206 participants was .95, .93, .97, .9 for the four factors.

The weights obtained for the four factors were multiplied onto the original 33 behavioural markers (z-scored) to construct four summary or latent behaviours per participant. These were summarized and are referred to throughout as ‘social and life satisfaction’, ‘negative emotions’, ‘sleep’ and ‘anger & rejection’.

### Regression analyses to identify the most predictive connections

A regression approach was used to identify the most predictive connections, separately for each of the four latent behaviours. The data to be predicted, **y**, was in each case a 200×1 vector describing the true latent behaviour for each participant. The matrix of potential predictors **X** was a matrix with 200 × 196 resting-state functional coupling (FC) values for each participant and the 196 connections described above (7 amygdala nuclei × 28 ROIs). Outlier participants were conservatively rejected based on their individual FC values if more than 10% of their FC values across all connections deviated more than 3.5 standard deviations from the mean across participants. This identified five participants as outliers and all analyses were performed on the remaining 195. Next, confounds were regressed out of the data as in ^101^. Confounds included (1) acquisition reconstruction software version; (2) summary statistic quantifying average head motion; (3) weight; (4) height; (5) blood pressure – systolic; (6) blood pressure – diastolic; (7) haemoglobin A1C in blood; (8) cube-root of total brain volume; cube-root of total intracranial volume. As described in detail in ^101^, in addition to these nine confounds, eight additional confounds included the demeaned and squared measures 2-9 to account for potential nonlinear confound effects. A total of 17 confounds were thus regressed out of the matrix X. Both y and X were z-scored.

For generating the plots in Fig 4, we estimated k=10,000 regression models using 10-fold cross-validation. For each model, we selected a random subset of five out of the total of 196 potential connections as predictor variables. We also generated a new cross-validation (CV) set in each iteration, with the additional constraint of keeping siblings together – i.e. all members of the same family were allocated together to the training set or to the test set. In each CV-fold, the goodness-of-fit was determined as the correlation (Pearson’s *r*) between true latent behaviour and the out-of-sample model-predicted behaviour obtained using the subset of five connections. For each model iteration, we saved the contributing connections and the average *r* across the 10 folds. The overall contribution of each connection (**Fig 4A-C**) was then determined across all 10,000 iterations as the average difference in *r* value between all iterations that did and all iterations that did not include the connection in the model. The distribution of these contribution values is shown across connections (Fig 4C), and we also report the histogram of raw *r* values from all 10,000 model iterations (Fig 4D).

It is worth highlighting some of the features of this procedure that explain its suitability for analysing our data. Including all connections in one large regression model was not feasible due to the large number of regressors and existing correlations between them. Our approach allowed us to identify two similar connections (e.g. NAc-LaV and NAc-LaD for sleep) as important because these two would only seldom be included simultaneously, by chance, in a regression model with five randomly selected connections. Using our approach, a smaller number of connections, e.g. considering each connection individually or only including two at a time in each sub-model, would over-estimate some contributions because shared features are assigned to each. On the contrary, larger sub-models with e.g. 20 or 30 connections would underestimate the predictors’ contributions. Importantly, we verified that our main conclusions were robust to choices of model size ranging from 2 to 5 to 10 connections (**Supplementary Figure 4**).

Fitting a large number of k=10,000 regression models allowed robust estimates of each connection’s contribution because with large enough k, contribution estimates converge (**Supplementary Fig 6D**). If we had estimated only k=200 models, for example, we would have, on average, estimated each connection’s contribution five times (200/196*5) and an average of five numbers would have been our final contribution estimate *rDiff*. By fitting 10,000 models, we estimated each connection 10,000/196 * 5 = ∼255 times, leading to a more robust estimate. At k=10,000 iterations, the estimated contribution *rDiff* changed very little with slight increases or decreases in the number of iterations: going from k=8,000 to k=10,000 iterations on average changed *rDiff* by .0058, going from k=10,000 to k=15,000 by .0055, indicating convergence (**Supplementary Fig 6D**). There was no risk of overfitting because all predictions were done out-of-sample.

To test whether contributions of individual connections were better than predicted by chance, given the level of noise present in brain connections with the amygdala and given our number of connections, we generated two versions of a null distribution (**Supplementary Fig 6A**) by instead predicting the vector **y** containing the latent behaviours using n=1,000 random sets of connections between all amygdala nuclei and 28 ROIs. In each of the n=1,000 iterations, we included all 196 connections between the seven amygdala nuclei and 28 randomly chosen ROIs (28*7=196; “amy-to-rnd”), thus matching the total number of connections with our main analysis of interest. We allowed connections from the original amygdala nuclei to *any* cortical region except our set of *a priori* ROIs. For each of the 1,000 iterations with random connections, we ran 1,000 sub-models with different sets of five connections and different CV-partitions, as above. We then extracted the distribution of the top connection to obtain p-values corrected for multiple comparisons, and of all connections to obtain uncorrected p-values (for illustration see **Supplementary Fig 6A**). To do this, we calculated the cumulative distribution function of the corrected and uncorrected distributions, which was used to generate the p-thresholds corresponding to FWE-corrected p<.05, and uncorrected p<0.001, respectively (denoted by black and grey asterisks in Fig 5A, respectively). For the illustration of the strongest connections in the fingerprints in Fig 6, we calculated the average regression coefficients for each connection to show the strength and sign of their influence on predicting the latent behaviour (shown as numbers on the connections in Fig 6). Scatterplots were produced for visual illustration of the strength of predictive power achieved with only the top connection of each latent behaviour (Fig 5C).

The above distributions were generated based on connections between the amygdala and randomly chosen other regions. As a result, they were likely conservative because we believe the amygdala itself carries importance for predicting variation in mental health and there might be other relevant connections with regions apart from the ones we specified *a priori*. In a second control, we tested whether contributions of our amygdala-to-ROI connections were superior to those obtained from connections with other subcortical regions. In each of n=1000 iterations, we chose seven random seeds of the same size as the amygdala nuclei, placed anywhere in HCP’s subcortical volume (containing NAc, brainstem, caudate, cerebellum, diencephalon, hippocampus, pallidum, putamen & thalamus). By using a subcortical seed and real brain connections, we matched the level of noise present in subcortical structures to our original analysis. For each of these n=1000 random subcortical seeds, we randomly chose 28 ROIs from anywhere in cortex, including our *a priori* ROIs (“subc-to-rnd”). This resulted in n=1000 hubs which closely matched the structure of our brain connections of interest. For each one, we performed out-of-sample estimations of the contribution of each connection, as above. Again, we generated null distributions by remembering the contribution *rDiff* of the top connection, or all connections, resulting in FWE-corrected and uncorrected *p*-values, respectively.

### Controlling for amygdala parcellation and ROI selection

To show that parcellating the amygdala yielded improvements in predictive power, we also repeated the regression procedure with only the connections from our ROIs to the entire amygdala instead of all individual nuclei (a total of 28 possible predictors). All figure panels related to connections with the whole amygdala, instead of its seven distinct nuclei, were generated using identical methods (**Fig 5B and Supplementary Fig 5**). Again, five connections went into each model and 1,000 iterations of models were generated using different CV-partitions. We generated separate null distributions for this analysis as before (k=1000, n=1000) and the *p*-thresholds for whole amygdala connections (Fig 5B) are relative to these new distributions which were generated based on (a) only the whole-amygdala to randomly selected ROI connections (“amy-to-rnd”) or (b) random subcortical (‘fake amygdala’) seeds to randomly selected ROI connections (“subc-to-rnd”). We also used the probability of each of our whole-amygdala to ROI connections, obtained from these uncorrected distributions, to generate adjusted alpha values against which we compared the corresponding nuclei-specific connections to the same ROI. This test established if parcellating the amygdala into nuclei helped us gain specificity in our predictions. For example, if the probability of the connection from p32pr to the whole amygdala, given the uncorrected amy-to-rnd distribution, is p=.02, we consider the parcellation to be a meaningful improvement if any of the nuclei-specific connections to p32pr are less than this adjusted alpha of .02. In this example, this was the case for the connection of p32pr with B (square symbol in Fig 5A; p<.001 given the nuclei-specific uncorrected distribution, and thus smaller than alpha of 0.02) but not any other nuclei-connections with p32pr. The same rationale can be used for both uncorrected control distributions (amy-to-rnd and subc-to-rnd) but the conclusions from both tests are virtually identical and therefore only reported for the former distribution (amy-to-rnd).

### Decoding latent behaviours

For decoding analyses, outlier rejection and regressing out of confounds was applied to the connectivity values as described above. For each latent behaviour, decoding was restricted to participants with scores in the top and bottom third and the behaviour was binarized. The predictors used by the decoder were the connections established as the top ten connections in each case above. We used a linear support vector machine (SVM, Matlab’s function fitclinear). The SVM was again trained on 90% and tested on the left-out 10% of values, with CV-folds respecting family structures, and this was repeated for all 10 folds. Prediction accuracy was computed as the area under the curve (AUC). P-values were derived from a histogram derived from bootstrapping (10,000 iterations) using behavioural and connectivity values that were shuffled between participants and respected family structure.

## Acknowledgements

Data were provided by the Human Connectome Project, WU-Minn Consortium (Principal Investigators: David Van Essen and Kamil Ugurbil; 1U54MH091657) funded by the 16 NIH Institutes and Centers that support the NIH Blueprint for Neuroscience Research; and by the McDonnell Center for Systems Neuroscience at Washington University. MCKF was funded by a Sir Henry Wellcome Fellowship (103184/Z/13/Z), MFSR was funded by an MRC grant (MR/P024955/1) and a Wellcome Senior Investigator Award (WT100973AIA).

## Author contributions

MCKF and MFSR designed the study, MCKF, DEAJ and MFSR conceived analyses, MCKF and DEAJ wrote analysis code, LV, YT and SS gave analysis advice, and all authors wrote the manuscript.

## Data availability statement

All data used in the present study are available for download from the Human Connectome Project (www.humanconnectome.org). Users must apply for access and agree to the HCP data use terms (for details see https://www.humanconnectome.org/study/hcp-young-adult/data-use-terms). Here we used both Open Access and Restricted data.

## Supplement

**Supplementary Note 1: Histogram of contributions**

We also examined histograms of, first, the contributions (**Fig 4C**) and, second, the underlying raw correlation coefficients across the *k*=10,000 regression models (**Fig 4D**). The distribution of the contributions across connections (**Fig 4C**) had a small tail to the right indicating a small number of predictive connections. Overall, the distributions were comparable across behaviours, although life satisfaction (blue) and sleep (green) had a slightly longer tail towards the right, indicating the existence of stronger predictors than for negative emotions and anger (95% confidence intervals: lifeSat: [-.044,.119], negEmot: [-.043,.102], sleep: [-.042,.101], anger: [-.036,.076]; consistent with Fig5A). The mode of the distributions was slightly to the left of zero, probably due to overfitting the training data when using non-predictive connections in the model which then generalize less well to the testing data (lifeSat: −.03, negEmot: −.03, sleep: −.02, anger: −.02). The raw correlation coefficients obtained across models (**Fig 5C**) were shifted slightly to the right of zero (mode: lifeSat: .02, negEmot: .02, sleep: .04, anger: .08), as expected if any of the connections meaningfully contribute to predict behaviour. The distribution for anger was shifted the most, indicating a larger number of connections, and thus a larger brain network, might help predict this behaviour.

**Supplementary Figure 1 – related to Figure 1.**
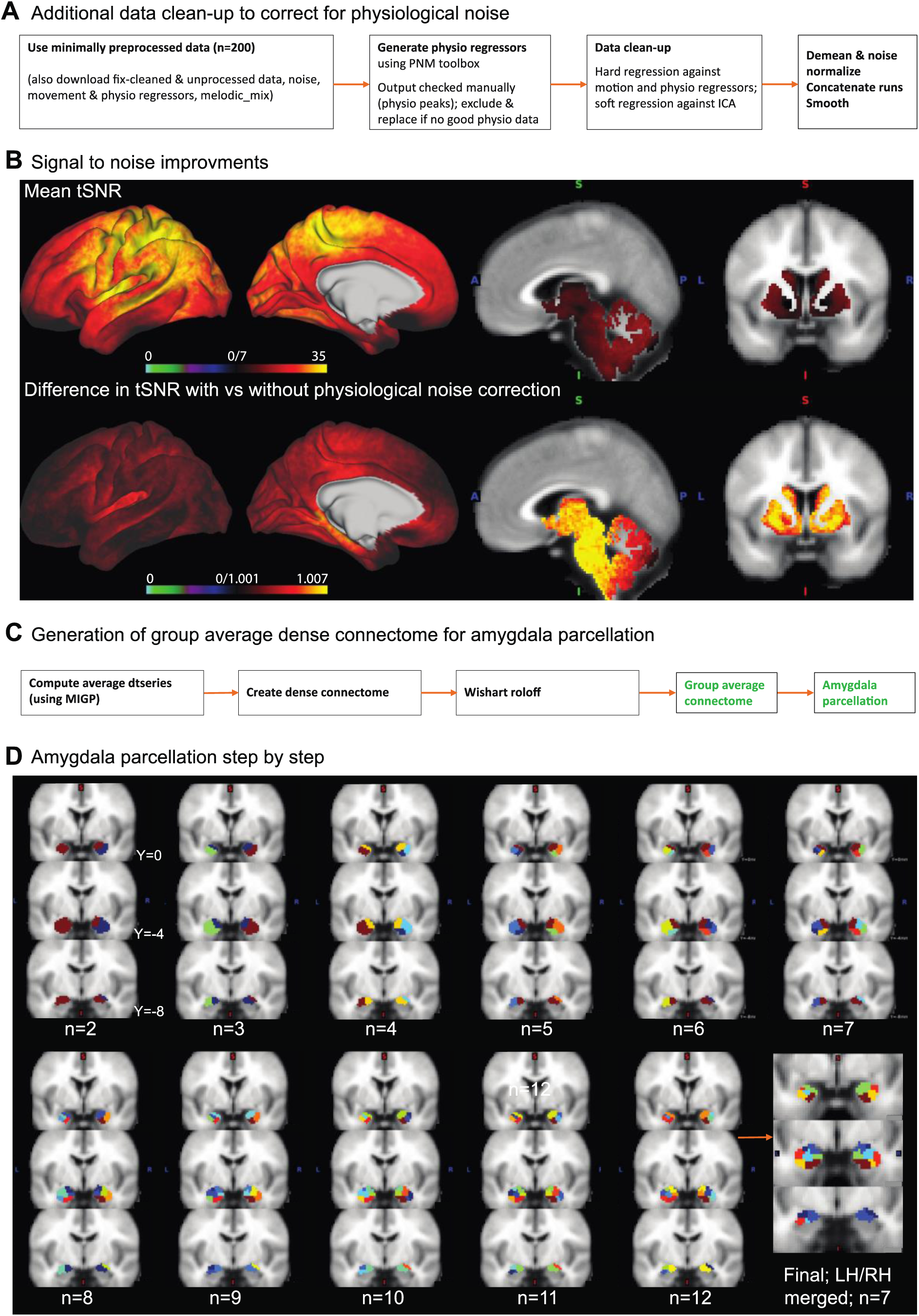
Preprocessing and hierarchical clustering pipelines. **A,** The minimally preprocessed HCP data was additionally corrected for physiological noise to improve the signal in temporal lobe and brainstem regions, the key areas for this study. All other data clean-up steps usually applied to generate fully preprocessed HCP data, specifically fix-denoising and motion correction, were applied at the same time. **B**, Illustration of the signal-to-noise improvements gained from this additional preprocessing step compared to standard full HCP preprocessing (in a subset of 100 participants). *Top*: Mean temporal signal to noise ratio (tSNR) obtained following our preprocessing pipeline; *Bottom*: Difference in tSNR between the preprocessing with and without physiological noise correction. The ratio of tSNRs (physio – noPhysio) / (physio + noPhysio) is illustrated. This shows tSNR gains in medial temporal lobe and medial prefrontal cortex but particularly subcortical and brainstem structures. **C**, Summary of the additional processing steps required to compute a group average connectome from the 200 individual concatenated resting-state fMRI (rs-fMRI) time-series. The group connectome, restricted to connectivity between amygdala voxels and the whole brain, formed the basis for the amygdala parcellation. **D**, Individual steps of the hierarchical clustering algorithm led to increasing subdivisions of the amygdala. All steps leading up to our final parcellation are shown. Hierarchical clustering was performed on absolute connectivity values. Note, for example, the central nuclei splitting off in step 9 (left) and 12 (right). The 12 cluster solution had five unique clusters in each hemisphere and two connected clusters (same color = same cluster). For subsequent analyses, the corresponding clusters in each hemisphere were joined, resulting in a total of seven clusters.

**Supplementary Figure 2 - related to Figure 3.**
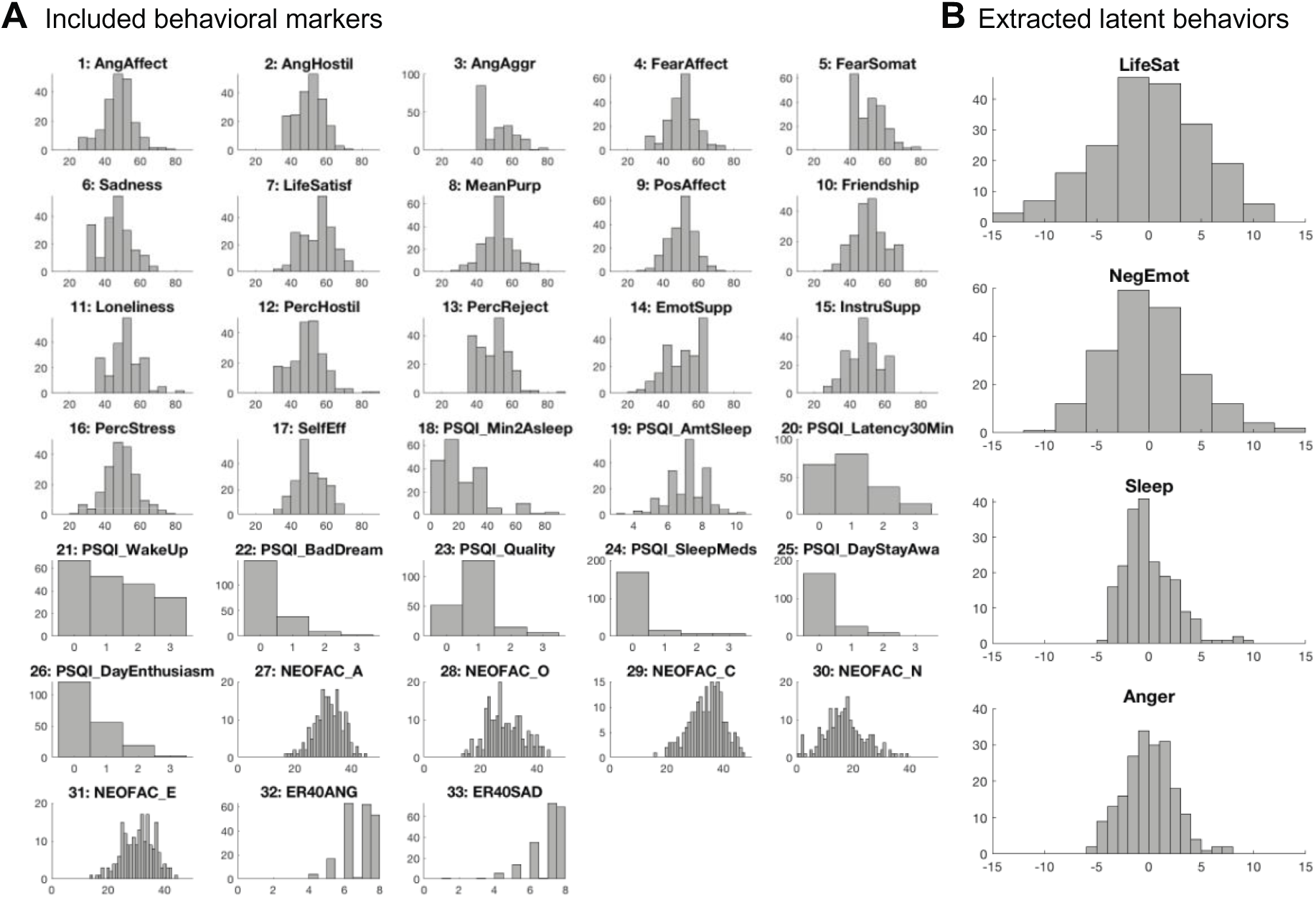
Distribution of behavioural scores and extracted latent behaviours. **A,** Distribution of all behavioural markers included in the factor analysis shown in Figure 3 across the 200 HCP participants. For a full description of each score see Table 1 and Methods. **B**, Distribution of the latent behaviours generated from the factor analysis.

**Supplementary Figure 3, related to Figure 4.**
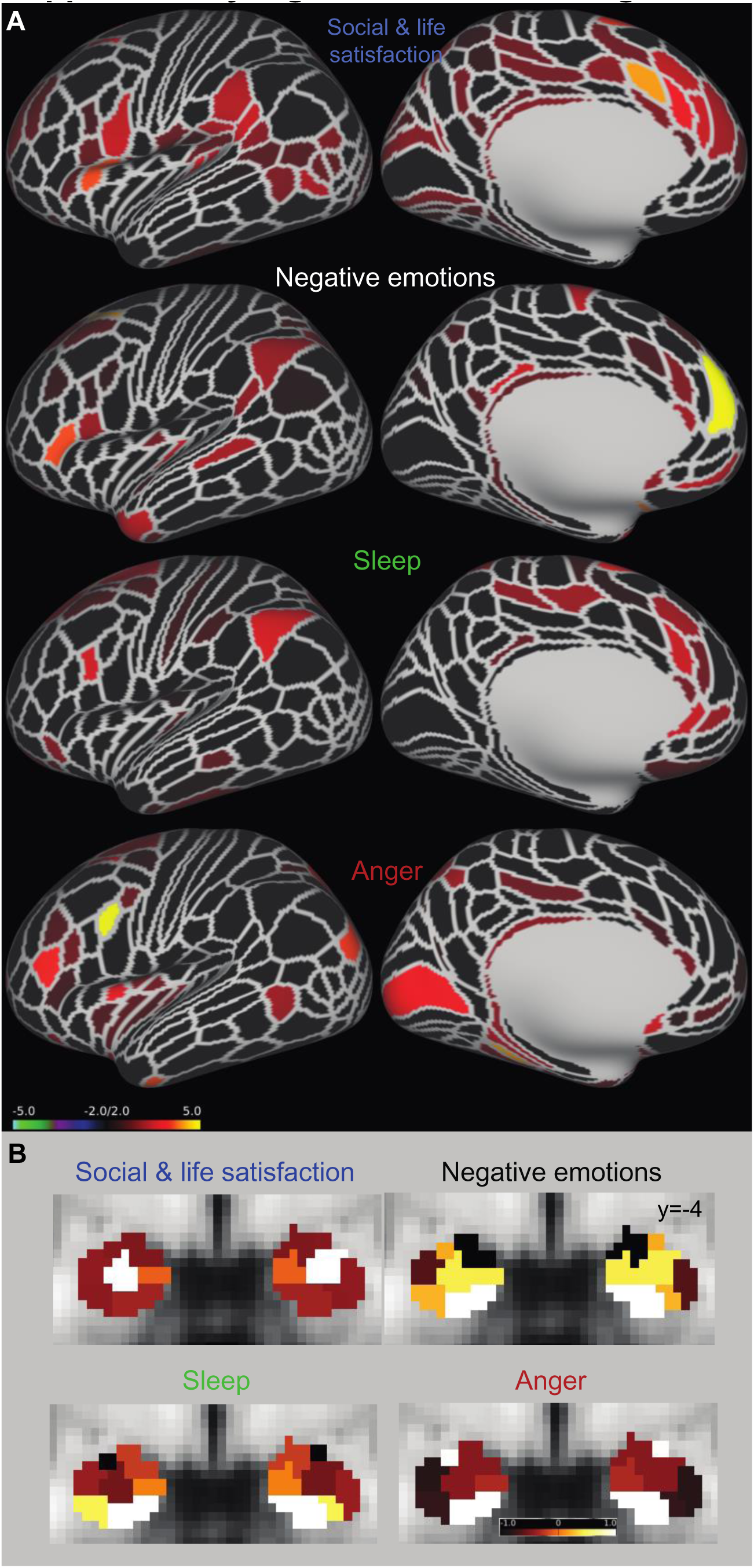
Maps of cortex and amygdala illustrate maximal contributions towards behavioural predictions. **A,** The distribution of *rDiff* values is shown for the entire cortex. *rDiff* values were z-scored across behaviours and connections. Each cortical parcel displays the *rDiff* value associated with the connection to the amygdala nucleus that was maximal for this cortical region. **B**, The contribution of the seven amygdala nuclei to each behaviour is shown. The values shown in the different colours summarize the contribution *rDiff* of connections with this nucleus for any instances when the connection with this nucleus was the top connection (out of all seven nuclei). Again, contribution values were z-scored across behaviours and connections.

**Supplementary Figure 4, related to Figure 5.**
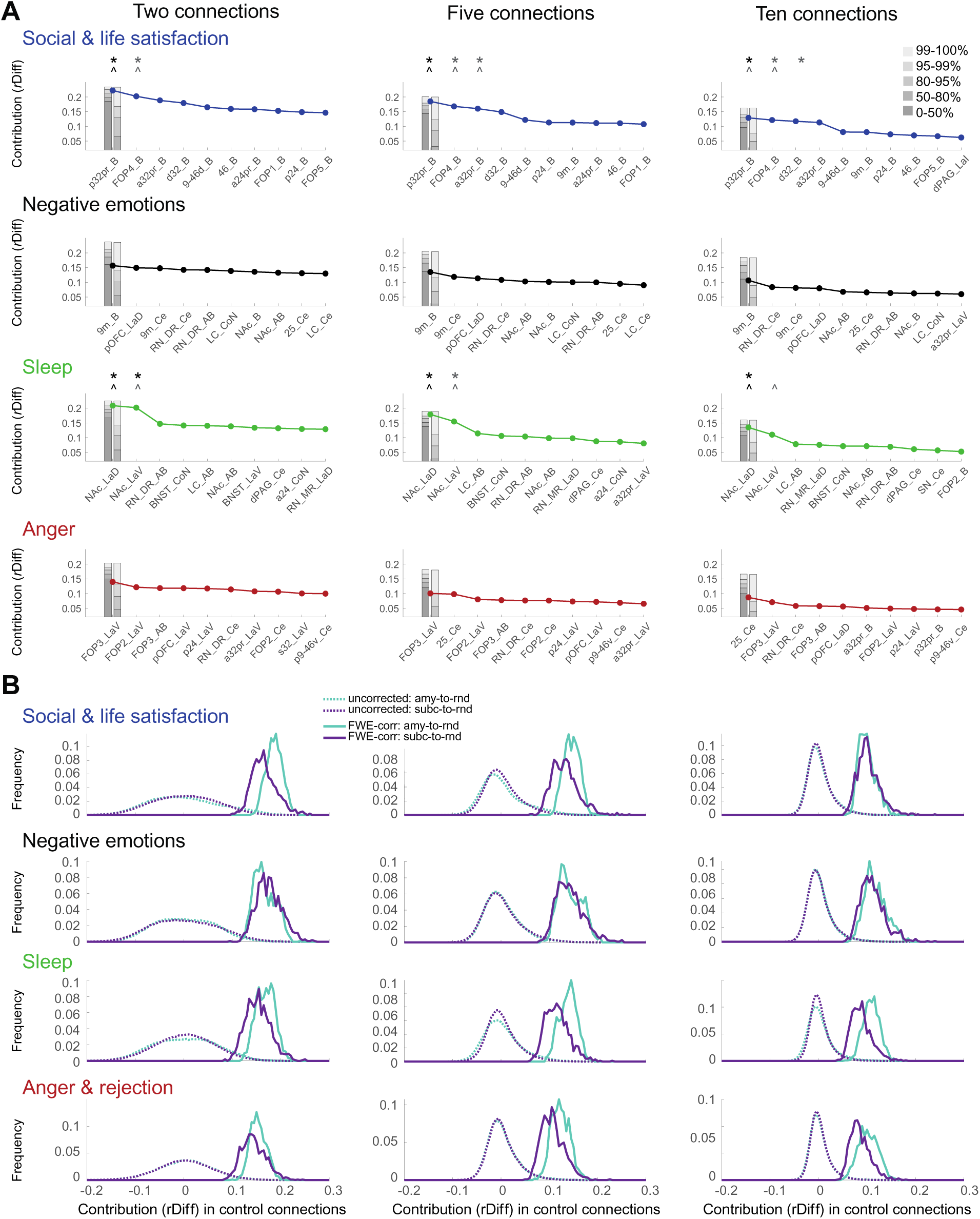
Predictions are robust to model size. **A,** Our main result in Figure 5A was obtained from 10,000 models with five randomly chosen connections each (reproduced here for comparison: middle column). We did not have enough data to optimize the number of connections included in each model as an additional hyperparameter. For transparency, here we therefore show the results for models involving only two (left column) or ten (right column) randomly selected connections in each of 10,000 model iterations. While small differences exist (such as the order of the top two connections flipping for Anger), none of the key results discussed in the paper are dependent on the selection of model size. As would be expected, an individual connection predicts slightly less variance (smaller *rDiff*), on average, when more regressors are included in the model (moving from left to right columns). **B,** However, this is taken into consideration in the generation of the respective control distributions which are used to establish significance (for more details, see Supplementary Figure 6).

**Supplementary Figure 5 – related to Figure 5.**
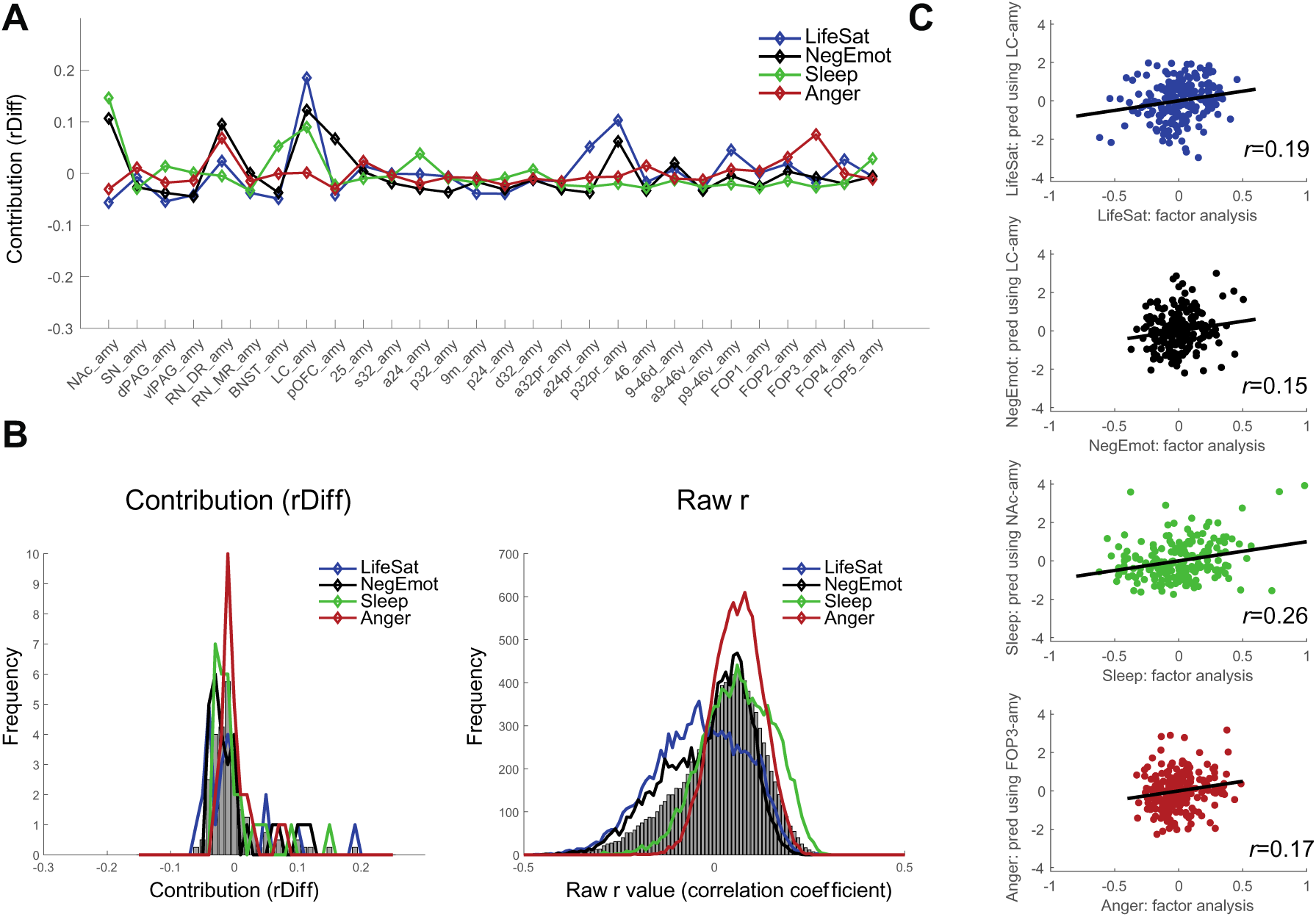
Mental well-being predictions benefit from parcellating the amygdala. To confirm that parcellating the amygdala into sub-nuclei increased our specificity for predicting mental well-being, we repeated the regression procedure using connections with the entire amygdala to the same 28 ROIs (see also **Figure 5B**). **A**, This highlighted LC-amy connectivity as important for predicting all latent behaviours except anger, NAc-amy connections for negative emotions and sleep and RN_DR-amy connections for negative emotions and anger, and thus primarily subcortical connections. Cortically, p32pr-amy connections were predictive of life satisfaction and negative emotions and FOP3-amy connection for anger. **B**, Histogram of contributions *rDiff* and raw *r* values are shown as in Figure 4C-D. **C**, The true behaviour obtained from the factor analysis is plotted against the behaviour predicted, in each case, using only the top connection with the whole amygdala. In summary, the anatomical specificity gained from parcellating the amygdala improved the prediction of mental well-being in the majority of cases (compare also **Figure 5A-B**).

**Supplementary Figure 6 – related to Figure 5.**
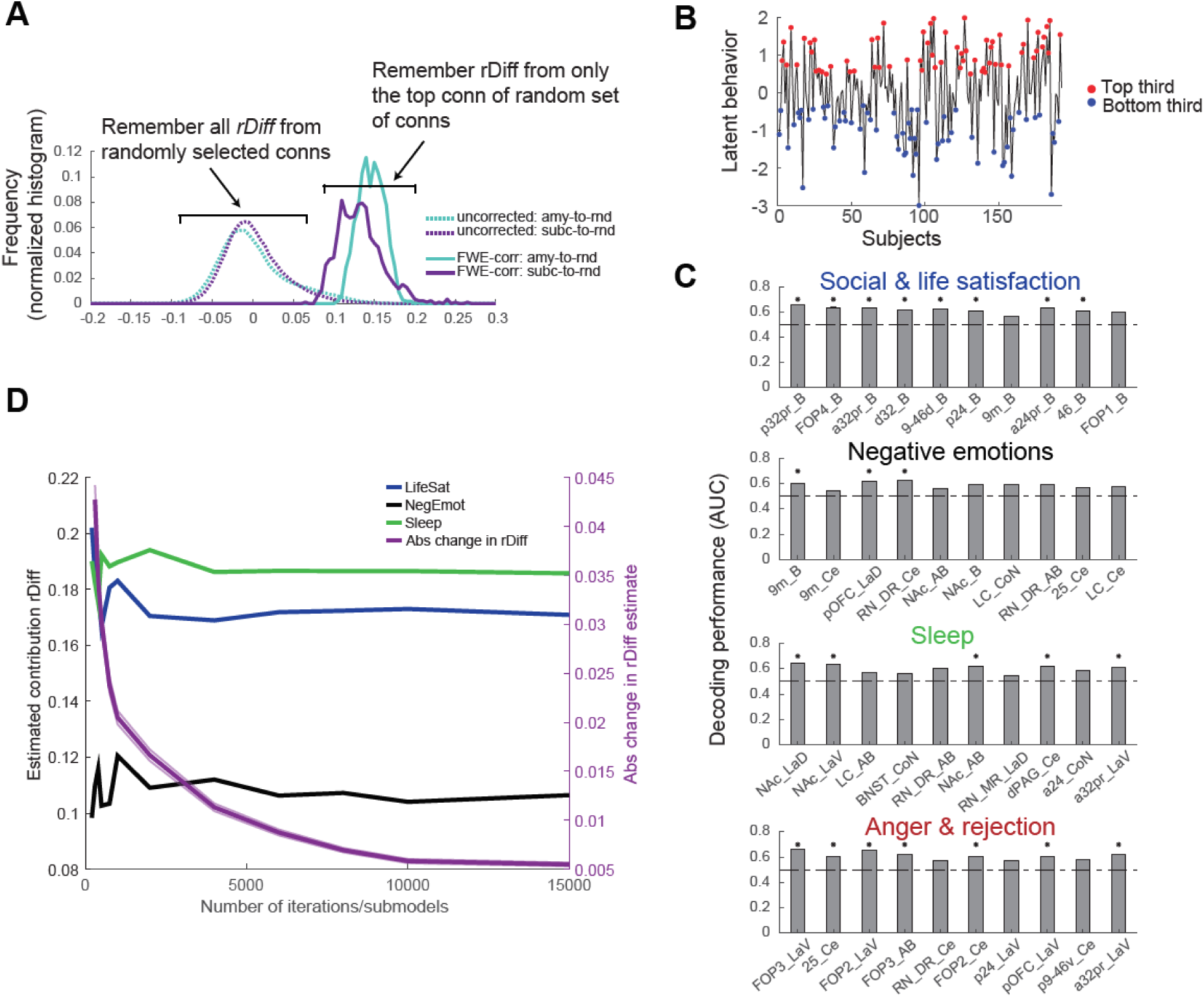
Illustration of statistical tests and iterations to convergence. **A**, We used two control analyses based on randomly selected connections, in each case matching the number of connections from our original amygdala-to-ROI analysis (7×28=196). In one case, random amygdala-to-cortex hubs with 28 cortical regions were created (“amy-to-rnd”), in the second case, random subcortical seeds of the same size as the original amygdala nuclei were defined, and hubs with these seven ‘fake’ nuclei and 28 randomly chosen cortical regions were constructed (“subc-to-rnd”). To generate uncorrected p-values, all 196 *rDiffs* were remembered in each of the 1000 random connection hubs and the resulting distributions are shown in the dashed lines and centred on 0. To correct for the number of connections tested (196), for each of the 1000 random hubs, we only remembered the top connection’s contribution. This led to the FWE-corrected distributions shown in the continuous line. In both cases, FWE-corrected and uncorrected p-values were generated using the cumulative distribution function (cdf) of the respective distributions. Distributions are shown exemplarily for life satisfaction here, but see Supplementary Figure 4B for all other behaviours. **B**, In an additional analysis, for comparison with other work that employs decoding techniques, we selected the top and bottom third of participants for each latent behaviour. This was done in order to maximize differences between our participants; note that our participants scored in a relatively narrow, sub-clinical range. Latent behavioural scores were binarized (1=high, 0=low). **C**, For the top 10 connections for each behaviour, the area under the curve (AUC) and thus decoding performance is shown. We were able to decode whether a participant was in the top or bottom third using multiple of the individual connections for all four latent behaviours. Significance was established using shuffled behavioural and connectivity values (see Methods and **Figure 5D**). **D,** The number of sub-models with five connections that were estimated to determine each connections’ contribution (*rDiff*) was set to k=10,000. To validate this choice, here we show (left y axis) the *rDiff* estimated for three somewhat relevant connections (d32-B for lifeSat, 25-Ce for NegEmot and NAc-LaV for sleep) as a function of the number of iterations/submodels that were estimated. This highlights that estimates of *rDiff* become more and more stable the more models are estimated. The right y axis shows the mean absolute difference in *rDiff* across all 196 connections that is seen between two subsequent choices of k. This shows that after about 8,000 iterations, estimates of *rDiff* hardly change, and that at 10,000 iterations, these estimates are robust and have converged.

